# Histone H2A variant H2A.B is enriched in transcriptionally active HSV-1 lytic chromatin

**DOI:** 10.1101/2023.12.22.573075

**Authors:** Esteban Flores, Sarah M. Saddoris, Arryn K Owens, Rebecca Gibeault, Daniel P. Depledge, Luis M. Schang

## Abstract

Herpes simplex virus 1 (HSV-1) transcription is restricted in latently infected neurons and the genomes are in mostly silenced chromatin, whereas all viral genes are transcribed in lytically infected cells, in which the genomes are dynamically chromatinized. Epigenetic regulation modulates HSV-1 transcription during lytic, latent, and reactivating infections, but the precise mechanisms are not fully defined. Nucleosomes are dynamic; they slide, breathe, assemble and disassemble. We and others have proposed that the most dynamic HSV-1 chromatin is transcriptionally competent whereas the least dynamic is silenced. However, the mechanisms yielding the unusually dynamic viral chromatin remain unknown. Histone variants affect nucleosome dynamics. The dynamics of H2A, H2A.X and macroH2A were enhanced in infected cells, whereas those of H2A.B uniquely decreased. We constructed stably transduced cells expressing tagged histone H2A, H2A.B, macroH2A, or H2B, which assembles the H2A/H2B nucleosome dimers with all H2A variants. All H2A variants, ectopic, and endogenous H2B, were assembled into HSV-1 chromatin evenly throughout the genome, but canonical H2A was relatively depleted from the viral chromatin whereas H2A.B was enriched in the most dynamic viral chromatin. When viral transcription was restricted, H2A.B became as depleted from the viral chromatin through the entire genome as H2A. We propose that lytic HSV-1 nucleosomes are enriched in the dynamic variant H2A.B/H2B dimers to promote HSV-1 chromatin dynamics and transcriptional competency, and conclude that the dynamics of HSV-1 chromatin are determined in part by the H2A variants.

**Importance:** HSV-1 transcription is epigenetically regulated during latent and lytic infections, and epigenetic inhibitors have been proposed as potential antiviral drugs to modulate latency and reactivation. However, the detailed mechanisms of regulation of HSV-1 transcription by epigenetics have not been fully characterized and may differ from those regulating cellular transcription. In particular, the lytic HSV-1 chromatin is unusually dynamic, whereas the latent silenced one is not, but the mechanisms resulting in the unique dynamics of the lytic chromatin remain unknown. Here we identify the enrichment on the highly dynamic histone 2A variant H2A in the most dynamic viral chromatin, which provides a mechanistic understanding for its unique dynamics. Future work to identify the mechanisms of enrichment in H2A.B on the viral chromatin may identify novel druggable epigenetic regulators that modulate HSV-1 latency and reactivation.

## Introduction

Herpes simplex virus 1 (HSV-1) is a nuclear replicating double-stranded DNA (dsDNA) virus. It establishes lytic infections during which all viral genes are transcribed, typically in mucosal epithelial cells, and latent infections, in which transcription is restricted, in sensory neurons.

The HSV-1 genomes are atypically chromatinized during lytic infections, but the roles of this chromatinization are still not entirely clear. HSV-1 DNA is transcribed by RNAPII, which normally transcribes chromatinized templates. Depending on the chromatinization state, however, chromatinization may also be a cellular defense mechanism to silence viral gene expression (1–3). To counteract this silencing, HSV-1 would have evolved mechanisms to increase the dynamics of HSV-1 chromatin. Consistently, all three HSV-1 transcription activators affect chromatin stability or dynamics. VP16 reduces the levels of total and acetylated H3 stably associated with HSV-1 immediate-early (IE) promoters (4); recruits histone acetyl transferases (GCN5, PCAF and CBP/p300) and chromatin remodeler SWI/SNF complexes to the IE promoters (5–7) and the histone demethylases LSD1 and JMJDs to decrease H3K9me3 and H3K27me3 modifications on HSV-1 chromatin (8–10); promotes Set1 and MLL1 mediated H3K4me3 (11, 12); and causes large-scale chromatin unfolding (13). ICP0 reduces the levels of total and acetylated H3 stably associated with early promoters (14); targets centromeric histone variant CenH3 for degradation (15–18); interacts with a regulator of the histone acetyltransferase CLOCK (19); disrupts the function of the histone H3.3 chaperone hDaxx1 (20, 21); decreases H3K9me3 and H3K27me3 accumulation of viral genes (22–24); counteracts ATRX-dependent nucleosome loading to viral genomes (25); disrupts the recruitment of hDaxx and HIRA to viral genomes, thus decreasing H3K9me3 and H3K27me3 (21, 26); displaces HDAC 1/2 from the REST/CoREST repressor complex, preventing deacetylation of viral chromatin (27, 28); promotes histone acetylation on viral chromatin (14); and promotes removal of silencing marks from HSV-1 chromatin (23). ICP4 interacts with the chromatin remodelers NuRD and INO80, the BRM and BRG-1 of the SWI/SNF complexes (29), and the histone acetyltransferase CLOCK (30). It also enhances histone dynamics, even in the absence of HSV-1 DNA or other HSV-1 proteins (31).

The basic unit of chromatin is the nucleosome, consisting of two core histone H2A-H2B dimers and one H3-H4 tetramer wrapped in 147 base pairs of dsDNA. Chromatin is dynamic. Histones disassemble from nucleosomes, diffuse through the nucleus, bound by chaperones, and reassemble nucleosomes at different sites (32–35). Often, cellular genes assembled in more dynamic nucleosomes are transcribed to higher levels than those assembled in less dynamic ones (36, 37).

Histones are conserved across eukaryotes. Yet, each histone is typically encoded in multiple genes, which often encode slightly different proteins and map to different chromosomes, (38, 39). H4 and H2B are the least divergent histones, with just one somatic variant including two isoforms of histone H4 (40–42) or one somatic variant including 13 isoforms, with no known functional differences, of H2B (43, 44).

H3 and H2A have evolved several functionally distinct somatic variants, with H2A being the most divergent core histone. Histone H3 encodes at least 3 functionally distinct variants, canonical H3.1/H3.2, which has three isoforms (42), H3.3, and CenH3. The functionally different H3.3 differs from H3.1/H3.2 by only five and four residues, respectively (43). Histone H2A has five functionally distinct variants in humans, H2A, H2A.X, H2A.Z, macroH2A, and H2A.B with eleven, one, three, three, and two isoforms, respectively (43, 45, 46).

H2A.B (H2A.Bbd type 1) was originally identified by its characteristic exclusion from the Barr body (47). H2A.B has only 48 % sequence homology with H2A (type 2A), lacking the heavily post-transcriptionally modified N-terminal lysine residues, as well as 15 residues at the C-terminus. H2A.B also lacks several of the lysine residues that contribute to the H2A “acidic patch”, which is critical for the inter-nucleosome interactions that stabilize both the nucleosome and higher order chromatin structures (48, 49). The H2A.B variant is highly expressed in testis, but it is then expressed to relatively higher levels in skin and less so in dorsal root and trigeminal ganglia than in all other tissues, in which is only minimally expressed-if at all (50). H2A.B expression in cultured somatic cells is inducible, being expressed to high levels in many cancer cells (51). Viral gene expression in primary human fibroblasts infected at a low multiplicity with HSV-1 was detected at 5 hpi specifically in the cells expressing the highest levels of H2A.B mRNA (52), and H2A.B mRNA levels increase in human neuronal cells infected in culture with any of the three HSV-1 strains tested (53).

The histone fold of macroH2A has 60 % sequence similarity to canonical H2A but its macro C-terminal domain is unique, as is its size of 42 kDa (54–56). macroH2A is enriched in the inactive X chromosome, the Barr body (54, 55). Nucleosomes assembled with macroH2A are less dynamic than those assembled with H2A (57–59), and genes assembled in nucleosomes enriched in macroH2A are generally silenced (60), possibly by restricting access of DNA to transcription proteins (61). H2A.X is the most conserved with H2A, having 91 % sequence similarity and differing mostly by a 13 amino acid long C-terminal tail. Phosphorylated H2A.X (γH2A.X) marks sites of DNA damage. H2A.Z has 59 % sequence similarity to canonical H2A (62), and it is enriched in nucleosomes at transcription start sites (60).

Unlike the canonical histones, which are synthesized mostly during S phase and are assembled in nucleosomes mostly via DNA replication-dependent mechanisms, most variant histones are synthesized throughout the cell cycle and assembled in nucleosomes via DNA replication-independent mechanisms. The replacement of canonical histone with their variants often modifies the stability of the nucleosomes.

H2A variants alter nucleosome dynamics. H2A.X assembles more dynamic nucleosomes than canonical H2A in vitro but is nonetheless less dynamic in cells (63, 64). H2A.Z assembles more dynamic nucleosomes than H2A (60, 65–68), particularly when assembled in nucleosomes containing H3.3 (66, 69, 70). macroH2A assembles less dynamic nucleosomes than canonical H2A (57, 71). H2A.B assembles the most dynamic nucleosomes of any H2A variant, being the most dynamic core histone (72, 73), and mimicking the N-or C-terminal tails deletions of H2A.B in canonical H2A increases its dynamics (32).

HSV-1 DNA is in highly dynamic chromatin during lytic infections and epigenetic regulation affects HSV-1 transcription (2, 3, 22, 74–80). However, the mechanisms yielding the highly unusually dynamic transcriptionally competent viral chromatin remain incompletely understood. All canonical core histones interact with several HSV-1 loci (81). As evaluated by chromatin immunoprecipitation (ChIP) efficiency, however, the interactions between histones and HSV-1 DNA appear to be weaker, sparser, or more dynamic than those with cellular DNA (6, 81–85). Consistently, there is an apparent depletion of histones in the herpes nuclear domains (HND), including the pre-replication and replication compartments. This apparent depletion may indicate that there are less histones in the HND, or reflect that histones are more dynamic in the HND than in the rest of the nucleus and thus spend less time in them (31). Nuclease protection assays of lytic HSV-1 chromatin result in heterogeneously sized DNA fragments that migrate as a smear in agarose gels and a main protected band, which is broader than expected from canonical nucleosomes (86, 87). Interestingly, histone H2A.B is typically enriched in highly transcribed cellular loci and is assembled into highly dynamic, including nucleolar, chromatin. Nuclease protection of H2A.B containing nucleosomes also results in heterogeneously sized DNA fragments that migrate as a smear in agarose gels and a major protected band broader than expected from canonical nucleosomes (88).

Here we show that the dynamics of H2A.B are differentially altered in HSV-1 infected cells and that H2A, macroH2A, and H2A.B are differentially assembled into H2A/H2B containing nucleosomes in HSV-1 chromatin. Whereas H2A, H2A.X and macroH2A had increased dynamics in infected cells, as H1, H2B, H3.1, H3.3, and H4 do (31, 89–91), H2A.B dynamics decreased. This unique decrease could result from the presence of new binding sites provided by HSV-1 DNA in infected cells. Consistently, H2A.B was preferentially incorporated into the most accessible HSV-1 chromatin when HSV-1 genes of all kinetic classes are transcribed. The distribution of H2A.B paralleled that of H2B through the entire HSV-1 genome, indicating no enrichment of H2A.B/H2B containing nucleosomes on any particular HSV-1 locus. When transcription was restricted to the IE loci with cycloheximide, H2A.B was as relatively depleted from HSV-1 chromatin as H2A, but still evenly distributed through the entire genome, including the transcribed IE loci.

We propose that HSV-1 DNA is preferentially assembled in unstable nucleosomes containing H2A.B/H2B dimers when the genomes are transcriptionally competent. This preferential assembly of H2A.B in H2A/H2B dimers in HSV-1 DNA nucleosomes would sequester the minor variant H2A.B resulting in the decrease in overall H2A.B dynamics and the atypical highly dynamic HSV-1 chromatin and unique protection pattern of the HSV-1 DNA. The relative enrichment in the dynamic variant H2A.B would promote global HSV-1 transcriptional competency, but not directly activate transcription of any given gene.

## Results

### EGFP-H2A.X, EGFP-H2A.B, and EGFP-macroH2A are incorporated into chromatin

Transiently expressed EGFP-H2A -H2A.X, -H2A.B, or -macroH2A localized to the nucleus with a granular discrete distribution, which is consistent with their incorporation into chromatin (Fig 1 A). Nuclei expressing EGFP-H2A.X, -H2A.B, or -macroH2A had well delimited bleached regions immediately after photobleaching still visible 100 seconds later, indicating their partial immobilization by incorporation into chromatin. Consistently with H2A.X assembling nucleosomes in vivo about as dynamic as those assembled with canonical H2A (63, 64), the average normalized fluorescence for EGFP-H2A dropped to approximately 15 % at 1 second, and recovered to 30 % in 100 seconds, consistent with previous experiments (32) (Fig 1 B, blue line), whereas the average normalized fluorescence for EGFP-H2A.X dropped to approximately 22 % at 1 second and recovered to 29 % in 100 seconds (Fig 1 B, blue line). Consistently with macroH2A assembling less dynamic nucleosomes than H2A, EGFP-macroH2A was less dynamic than EGFP-H2A. The fluorescence intensity of the bleached region was not noticeably different between nuclei expressing EGFP-macroH2A or -H2A at 1 second after photobleaching (Fig 1 A, B, blue line), but the recovery was faster for EGFP-H2A in that EGFP-macroH2A had recovered only 20 % of fluorescence in 100 seconds (Fig 1 B, blue line).

**Figure 1.**
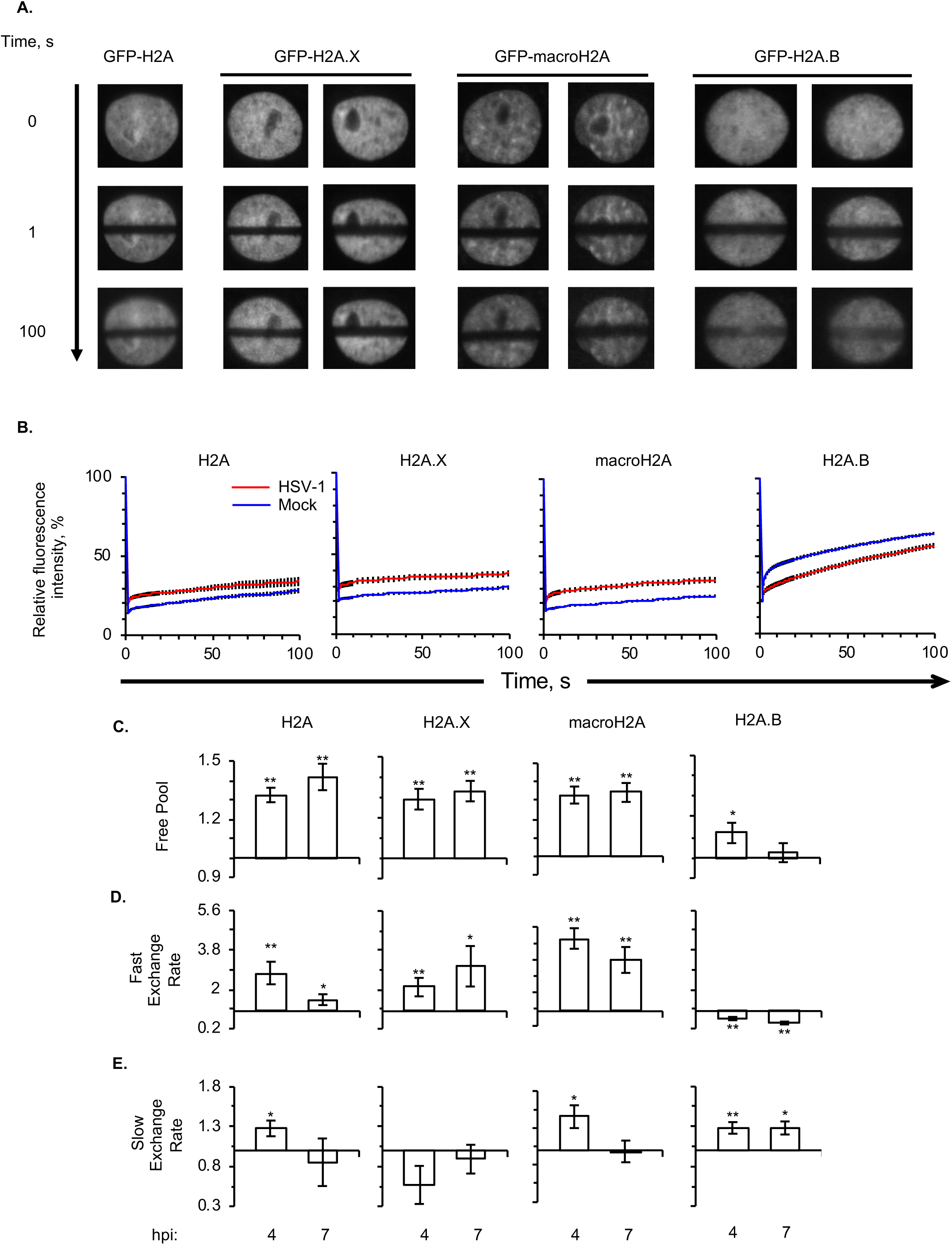
The dynamics of H2A, H2A.X, and macroH2A are enhanced in HSV-1 infected cells, whereas those of H2A.B decrease. A) Representative fluorescent micrograph images of Vero cells expressing EGFP-H2A, -H2A.X, -H2A.B, or -macroH2A, 1 and 100 seconds after photobleaching the equatorial region. B, C, D, E) Vero cells were transfected with a plasmid expressing EGFP-H2A, -H2A.X, -H2A.B, or macroH2A. Transfected cells were mock-infected or infected with HSV-1 moi 5. Dynamics of EGFP-H2A, -H2A.X, -H2A.B, or macroH2A were evaluated 4 to 5 (4 hpi) or 7 to 8 (7 hpi) hours later by FRAP. B) Average fluorescence recovery curves for EGFP-H2A, -H2A.X, -H2A.B, or macroH2A at 7 hpi in HSV-1- (red) or mock-infected (blue) cells. Average ± SEM. n ≥ 15 cells from at least 3 independent experiments. C) Bar graphs showing the average levels of free EGFP-H2A, -H2A.X, -H2A.B, or macroH2A in HSV-1-relative to mock-infected cells at 4 or 7 hpi. D) Bar graphs showing the average fast exchange rates of EGFP-H2A, -H2A.X, -H2A.B, or macroH2A in HSV-1-relative to mock-infected cells at 4 or 7 hpi. E) Bar graphs showing the average slow exchange rates of EGFP-H2A, -H2A.X, -H2A.B, or macroH2A in HSV-1-relative to mock-infected cells at 4 or 7 hpi. Hpi: hours post infection. Average ± SEM. **, P < 0.01; *, P < 0.05. n ≥ 15 cells from at least 3 independent experiments.

EGFP-H2A.B was the most dynamic EGFP-H2A variant, consistent with H2A.B assembling the most dynamic nucleosomes. The fluorescence intensity of the bleached region recovered more in 100 seconds in the nuclei expressing EGFP-H2A.B than EGFP-H2A (Fig 1 A, B, blue line). The average normalized fluorescence of EGFP-H2A.B was greater than 20 % at 1 second and recovered to as much as 70 % in 100 seconds (Fig 1 B, blue line), consistently with previously published results (72).

The dynamics of H2A, H2A.X, and macroH2A all increased in infected cells, as previously shown for H2A and their partner H2B (31, 90). The fluorescence recovery of H2A, H2A.X, and macroH2A was faster in infected than mock-infected cells and their free pools and fast exchange rates increased at both 4 and 7 hpi (Fig 1 B, red and blue lines, C, D), consistent with previously published results obtained with H2A (31), H2B (90), H3 (91), and H4 (90). By contrast, the fluorescence recovery of H2A.B was slower in infected than mock-infected cells, with the fast exchange rate being uniquely slower at 4 or 7 hpi (Fig 1 D). The free pool of H2A.B slightly but significantly increased at 4 h, but less so than for the other H2A variants, and then it decreased at 7 hpi, in contrast to all other variants.

In summary, the overall dynamics of H2A.B decreased in HSV-1 infected cells, while its slow exchange rate increased. One possibility is that HSV-1 DNA provides novel binding sites for H2A.B and thus a larger fraction of H2A.B would spend more time assembled into chromatin, even if the nucleosomes are unstable, decreasing the free pool and the fast exchange rate while increasing the slow one.

### Flag-H2A, -H2A.B, -macroH2A, and -H2B are incorporated into chromatin and result in no major obvious impairment in replication or ability to support HSV-1 replication

No antibody recognizes the H2A.B variant, or discriminates H2A from all other variants. To evaluate whether the HSV-1 chromatin is enriched or depleted in different histone H2A variants, we generated HeLa cells stably expressing flag tagged H2A, H2A.B, macroH2A, or H2B. We selected H2A.B based on its unique properties, its relatively higher level of expressions in skin and dorsal root or trigeminal ganglia (50), its higher expression in cells supporting HSV-1 gene expression (52), induction in HSV-1 infected cells (53), and the previous results suggesting incorporation into viral chromatin (Fig 1 B). macroH2A was selected based on its dynamics, its association with stable latent HSV-1 chromatin (92), and its relatively lower expression levels in cells supporting HSV-1 gene expression (52). H2A.X was not included as its dynamics were affected in parallel to those of canonical H2A. We used human HeLa cells, which have been used to study histone H2A variants using similar approaches (60, 93) and are derived from cervical epithelial cells. We changed the tag from EGFP to the smaller 3xflag to minimize potential effects of a larger tag.

The constitutively expressed flag tagged histones were all assembled into chromatin (Fig 2 A), as the transiently expressed GFP tagged ones (Fig 1 A, (31, 72)), as evaluated by colocalization with DNA in discrete distribution and localization in mitotic and interphase cells. Except for H2A.B, all ectopically expressed flag tagged histones were in the most condensed chromatin throughout mitosis (Fig 2 A). In contrast, H2A.B appears to be excluded from the most condensed chromatin in cells from metaphase through anaphase and telophase, to be incorporated again at late telophase (Fig 2 A), as previously shown (72) and expected from a histone that assembles so dynamic nucleosomes (72, 73) (Fig 1). We conclude that the flag tag was appropriate to evaluate all histone variants with the same antibody.

**Figure 2.**
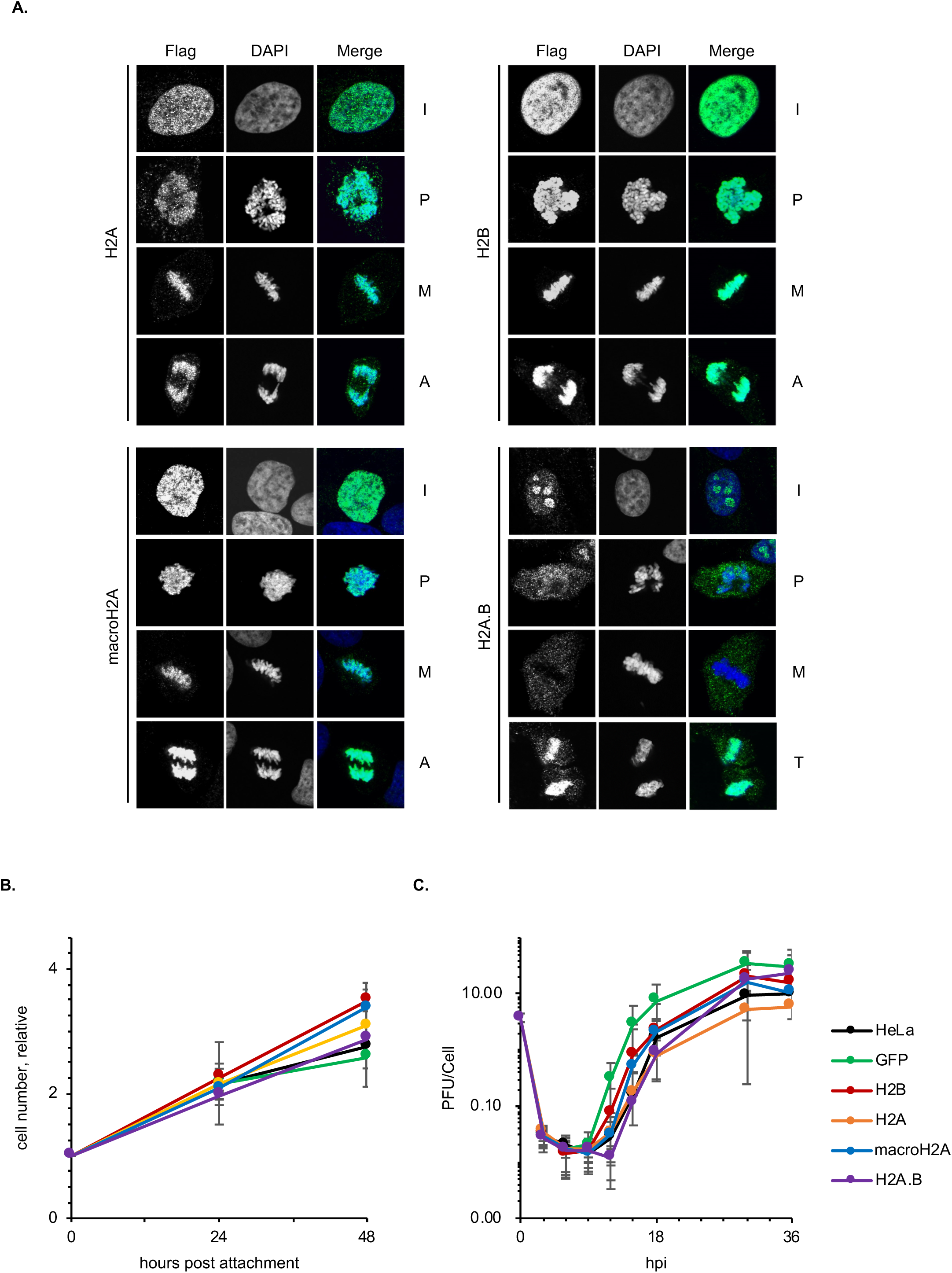
Constitutively expressed flag-H2A, flag-H2A.B, flag-macroH2A, and flag-H2B are incorporated into chromatin and their expression results in no major obvious differences in cell division or HSV-1 replication. A) Immunofluorescent micrographs of HeLa cells expressing flag tagged H2A, H2A.B, macroH2A, or H2B stained with Alexa Fluor 488 labeled anti-flag (green) or DAPI (blue). Cells at interphase, prophase, metaphase, or late telophase stages of the cell cycle, as evaluated by DAPI staining. Note that H2A.B shows disperse nuclear staining in some cells in interphase and that its location to condensed chromatin is highly infrequent, and most likely restricted to late telophase. I, interphase; P, prophase; M, metaphase; A, anaphase; T, late telophase. B) Relative cell number for each cell line during 48 h evaluated by CellTiter-Glo 2.0 kit; n = 3; Average ± SD. C) HSV-1 replication curves in the different cells. Hpi: hours post infection; n = 5; Average ± SD.

We assessed the effects of the tagged histones on cell doubling time. The doubling times for the cell populations expressing any flag histone, or EGFP, differed by less than four hours from that of non-transduced HeLa cells (Fig 2 B). We next evaluated HSV-1 replication kinetics in each cell population. Although there was some level of variation, all supported viral replication with similar lag times followed by the exponential increase in viral titers and ensuing plateau at similar levels (Fig 2 C). We concluded that the expression of flag tagged histones did not have any major obvious effects on cell doubling or HSV-1 replication and proceeded to evaluate their incorporation into HSV-1 chromatin.

### H2A.B is less depleted in the HND than H2A or macroH2A

To test whether canonical histone H2A and variants H2A.B and macroH2A may be differentially localized in the HND, we infected the cells expressing each flag-tagged histone variant and labelled replicating HSV-1 DNA with 5-ethynyl-2’-deoxyuridine and 5-ethynyl-2’-deoxycytidine for 2 hours starting at 6 hours after infection. Cells were fixed and processed via immunofluorescence and click chemistry, and visualized by confocal microscopy. Replicating viral DNA was preferentially labeled over cellular DNA, as evaluated by the large colocalization of the replicating DNA signal with ICP8, or immediately next to it (Fig 3 A). The relative localization of each flag-tagged histone variant and replicating viral DNA was then evaluated in comparison to their co-localization with replicating cellular DNA in uninfected cells. Whereas all variants co-localized with replicating cellular DNA, and as expected H2A.B co-localized the most (Fig 3 B) (94), H2A and macroH2A were depleted by more than half in the HND, while H2A.B was much less so (Fig 3 C, D). H2A.B preferentially localizes to replicating cellular DNA (94), and therefore its relative enrichment in the HND may result active ongoing HSV-1 DNA replication. Nonetheless, the differential localization to the HND may suggest that different variants may be differentially assembled into HSV-1 chromatin.

**Figure 3.**
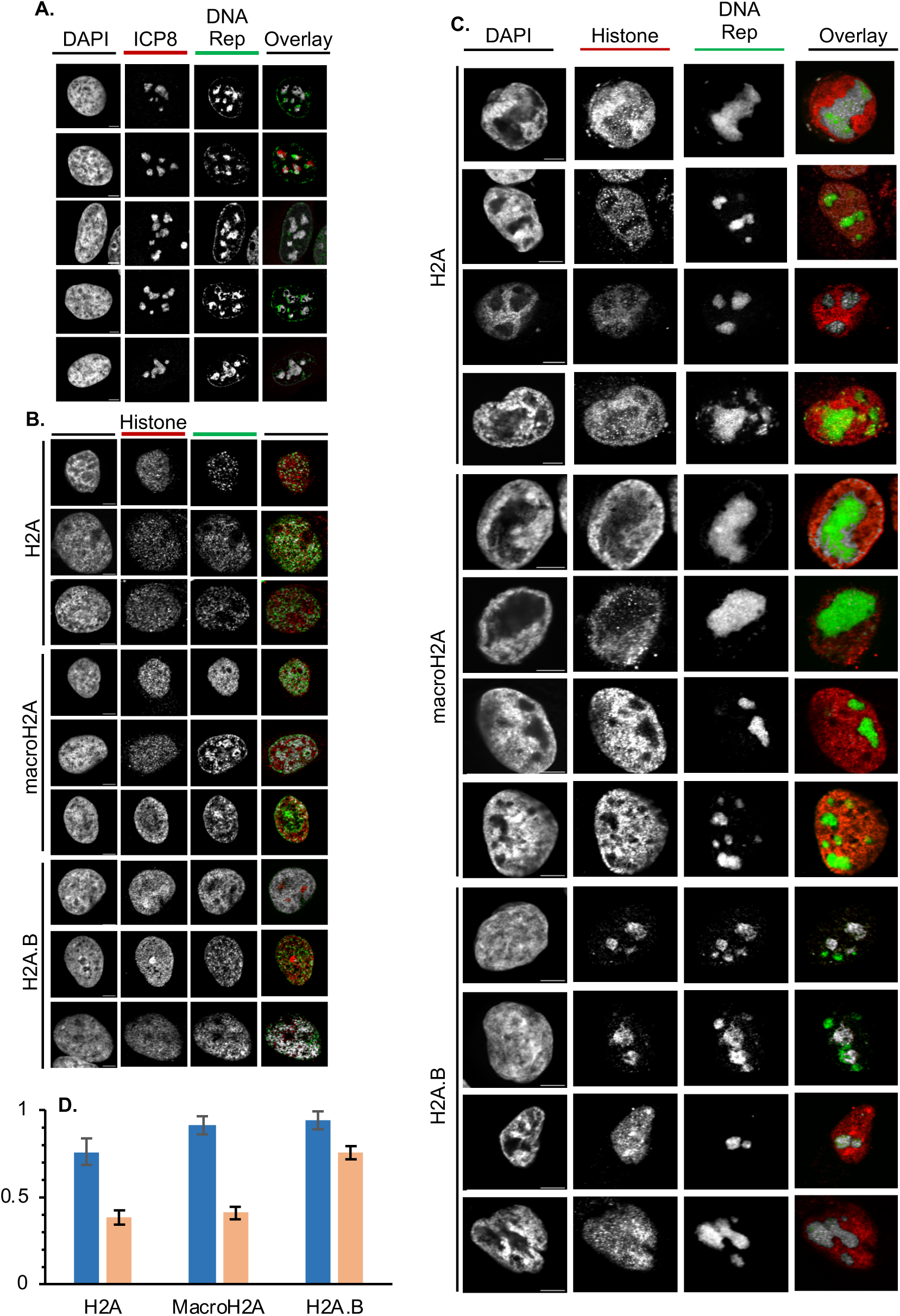
H2A.B is less depleted than H2A or macroH2A from the HND. Localization of replicating HSV-1 DNA relative to each histone variant of interest. Cells expressing each flag-tagged histone variant (B-D), nor not (A), were infected with HSV-1 (MOI=3; A, C, D), or not (B, D), for 8 hours. Replicating DNA was labelled at from 6 to 8 hpi with 5-ethynyl-2’-deoxyuridine and 5-ethynyl-2’-deoxycytidine, and its localization relative to ICP8 (A) or the indicated histone variant (B, C) was evaluated by immunofluorescence, click chemistry, and confocal microscopy. Single channels in black and white and overlays in color (red, ICP8 or histone; green, replicating DNA; grey, co-localization). C, Bar graph presenting the percentage of replicating DNA co-localizing with each histone H2A variant in uninfected or HSV-1 infected cells, avg±SEM. Scale bar = 5μm; n=5.

### All tested histone H2A variants are assembled into HSV-1 chromatin regardless of chromatin dynamics

To test whether ectopic expression of canonical H2A, H2A.B, macroH2A, or H2B affected HSV-1 chromatin dynamics, chromatin of cells expressing each flag tagged histone and infected with HSV-1 for 7 hours was serially digested with MCN followed by centrifugal fractionation. Fractionated chromatin was crosslinked in solution, specifically co-immunoprecipitated with anti-flag or -H2B antibodies, and DNA shorter than 570 bp (about tri-nucleosome length) was sequenced. Insert length frequency was analyzed for 50,000 cellular or viral alignments, except for the samples from the non-digested and non-fractionated chromatin of the H2A.B expressing cells co-immunoprecipitated with anti-flag antibody and the macroH2A cells co-immunoprecipitated with anti H2B antibody in the first biological repeat, and of the H2A.B expressing cells co-immunoprecipitated with H2B antibody in the second. For these three samples, subsampling was reduced to 12,000 cellular or viral alignments due to limiting number of viral reads.

HSV-1 DNA was quantitatively protected from MCN during serial digestion. HSV-1 DNA was protected on average across all flag-tagged cells 54 to 67 % as efficiently as the cellular DNA in each of the two biologically independent experiments, in the ranges of previous results (2, 75). When analyzing the cells expressing the different flag-tagged histones, HSV-1 DNA was consistently 1.5 or 1.3-fold better protected than cellular DNA in the flag tagged H2A.B expressing cells than in the flag tagged H2A or macroH2A expressing ones, respectively. HSV-1 DNA recovery in the cells expressing flag tagged H2A.B was 70 or 80 % as efficient that of the cellular DNA in each of the two individual experiments. We have shown that naked HSV-1 DNA added to chromatin from mock infected cells is not qualitatively protected in serial MCN digestion (2).

The protected cellular DNA fragments had a mean length of 206 bp and an average mode of 170.5 bp through all digested fractions (Table 1, 2, Cellular DNA, noIP), with a somewhat narrower distribution and more marked mode at 167 bp and a mean fragment length of 191 bp in the most accessible cellular chromatin (Table 1, 2, Cellular DNA, short, noIP, Fig 4). The distribution of the digested HSV-1 DNA fragments had a slightly longer fragment mean (213 bp) and mode (183 bp) and a less centered distribution than the cellular ones (Table 1, 2, HSV-1, noIP). These results are consistent with the previously described broader distribution of the protected DNA fragments in the viral chromatin (74). Like for the cellular DNA, the mean and mode of the HSV-1 DNA fragments in the most accessible chromatin were somewhat shorter than in the mid accessible or pelleted chromatin (200 and 172 bp, respectively, Table 1, 2, HSV-1, short, noIP). The modes of the most accessible viral and cellular chromatin are consistent with 147 bp protected by a core nucleosome plus about 20 bp long of cellular, or slightly longer (25 bp) of viral, DNA. This difference is consistent with the concept that the viral nucleosomes are more dynamic, sliding and protecting longer DNA during the restricted mild digestions, and inconsistent with the presence of mostly naked viral DNA, which should be more accessible and easily digested to smaller fragments than chromatinized cellular DNA.

**Table 1.** Median length of the DNA fragments protected from MCN digestion and immunoprecipitated with anti-flag or anti-H2B antibodies by fraction (averages for the cell lines expressing any flag tagged histone) or histone (averages of all the fractions for each cell line).

|  |  | HSV-1 |  |  | Cellular |  |  |
| --- | --- | --- | --- | --- | --- | --- | --- |
|  |  | noIP | Flag | H2B | noIP | Flag | H2B |
| Undigested<br>Unfractionated |  | 167.00 | 182.63 | 185.63 | 194.13 | 216.00 | 213.75 |
| Serially<br>digested<br>fraction | Short | 199.88 | 218.00 | 215.88 | 190.75 | 221.00 | 247.88 |
|  | Long | 226.13 | 242.13 | 239.63 | 207.75 | 247.00 | 244.63 |
|  | Pellet | 213.25 | 239.38 | 236.00 | 219.75 | 247.00 | 244.63 |
|  | *Mean* | *213.08* | *233.17* | *230.50* | *206.08* | *238.33* | *245.71* |
| Histone<br>H2A variant | H2A | 208.50 | 229.83 | 232.00 | 203.83 | 232.33 | 251.33 |
|  | H2A.B | 209.00 | 226.50 | 226.00 | 207.67 | 238.67 | 243.67 |
|  | macroH2A | 218.67 | 236.17 | 228.33 | 205.50 | 239.83 | 244.33 |
|  | H2B | 216.80 | 238.54 | 233.67 | 205.33 | 238.83 | 242.67 |

**Table 2.**
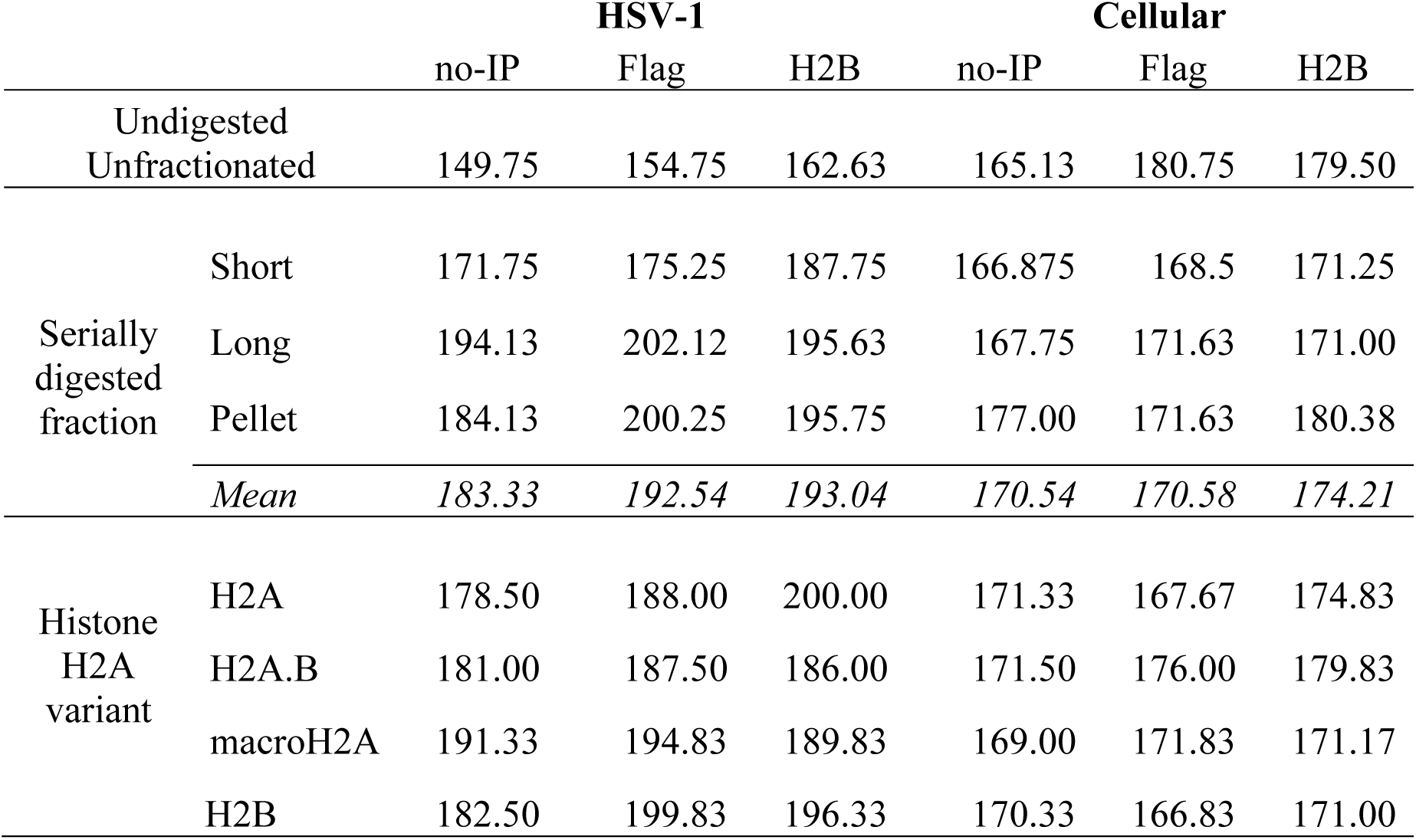
Mode of the length of the DNA fragments protected from MCN digestion and immunoprecipitated with anti-flag or anti-H2B antibodies by fraction (averages for the cell lines expressing any flag tagged histone) or histone (averages of all the fractions for each cell line).

**Figure 4.**
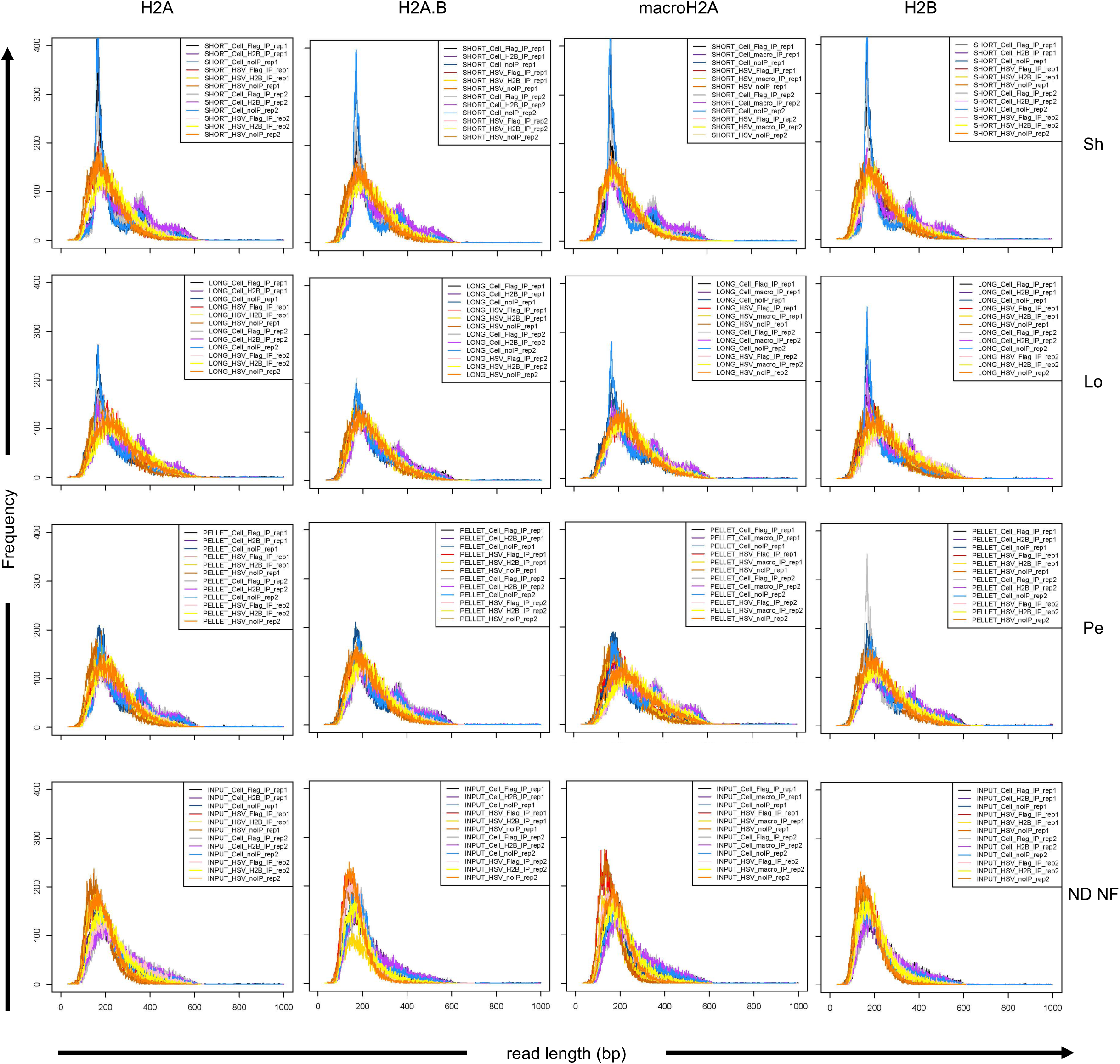
HSV-1 DNA is protected to, and co-immunoprecipitated with histone H2A and H2B in, fragments sizes consistent with nucleosome DNA. Chromatin of HeLa cells expressing each flag tagged histone and infected with HSV-1 for 7 hours were serially digested with MCN followed by fractionation. Fractionated chromatin was crosslinked in solution, co-immunoprecipitated or not with anti-flag or anti-H2B antibodies, and DNA reads shorter than 570 bp were analyzed. Fragment length distributions are shown with fragment length along the x-axis and the abundance count shown on the y-axis.. Black line, cellular chromatin IP with flag antibody biorepeat 1; Dark purple line, cellular chromatin IP with H2B antibody biorepeat 1; Dark blue line, cellular chromatin no IP biorepeat 1; Red line, HSV-1 chromatin IP with flag antibody biorepeat 1; Yellow line, HSV-1 chromatin IP with H2B antibody biorepeat 1; Orange line, HSV-1 chromatin no IP biorepeat 1; Gray line, cellular chromatin IP with flag antibody biorepeat 2; Purple line, cellular chromatin IP with H2B antibody biorepeat 2; Blue line, cellular chromatin no IP biorepeat 2; Pink line, HSV-1 chromatin IP with flag antibody biorepeat 2; Light yellow line, HSV-1 chromatin IP with H2B antibody biorepeat 2; Light orange line, HSV-1 chromatin no IP biorepeat 2; Sh, short accessible chromatin; Lo, long accessible chromatin; Pe, insoluble chromatin; ND NF, non-digested non-fractionated but sonicated chromatin. Biorepeat, biologically independent experiments.

To test whether the HSV-1 DNA protection patterns result from HSV-1 chromatin assembled with nucleosomes containing canonical H2A, H2A.B, or macroH2A in the H2A/H2B dimers, we analyzed the fragment size distribution of HSV-1 DNA co-immunoprecipitated with H2B or the H2A variants. The DNA fragments co-immunoprecipitated with any histone had slightly longer mean than the total protected DNA, 32 to 39.5 bp longer for cellular DNA co-immunoprecipitated with the flag or H2B antibodies (Table 3, Cellular [IP]-[noIP]), or 20 to 17.5 for the viral DNA co-immunoprecipitated with flag or H2B antibodies, respectively (Table 3, HSV-1 [IP]-[noIP], Fig 4). The modes were also slightly longer for HSV-1 DNA co-immunoprecipitated with flag or H2B antibodies, but only by 9.2 to 9.7 bp, respectively (Table 4, HSV-1, (IP-noIP)), indicating that the means were biased by the increased co-immunoprecipitation of longer DNA, likely in (longer) dynamic di- and tri-nucleosomes.

**Table 3.** Increase in median length of the DNA fragments protected from MCN digestion and immunoprecipitated with anti-flag or anti-H2B antibodies over the total DNA fragments protected from MCN digestion by fraction (averages for the cell lines expressing any flag tagged histone) or histone (averages of all the fractions for each cell line).

|  |  | **HSV-1**<br>(IP)-(noIP) |  | **Cellular**<br>(IP)-(noIP) |  |
| --- | --- | --- | --- | --- | --- |
|  |  | Flag | H2B | Flag | H2B |
| Undigested<br>Unfractionated |  | 15.63 | 18.63 | 21.88 | 19.63 |
| Serially<br>digested<br>fraction | Short | 18.13 | 16.00 | 30.25 | 57.13 |
|  | Long | 16.00 | 13.50 | 39.25 | 36.88 |
|  | Pellet | 26.13 | 22.75 | 27.25 | 24.88 |
|  | *Mean* | *20.08* | *17.42* | *32.25* | *39.63* |
| Histone<br>H2A<br>variant | H2A | 21.33 | 23.50 | 28.50 | 47.50 |
|  | H2A.B | 17.50 | 17.00 | 31.00 | 36.00 |
|  | macroH2A | 17.50 | 9.67 | 34.33 | 38.83 |
|  | H2B | 21.74 | 20.83 | 33.50 | 37.33 |

**Table 4.** Changes in mode length of the DNA fragments protected from MCN digestion and immunoprecipitated with anti-flag or anti-H2B antibodies over the total DNA fragments protected from MCN digestion by fraction (averages for the cell lines expressing any flag tagged histone) or histone (averages of all the fractions for each cell line).

|  |  | HSV-1<br>(IP)-(noIP) |  | Cellular<br>(IP)-(noIP) |  |
| --- | --- | --- | --- | --- | --- |
|  |  | Flag | H2B | Flag | H2B |
| Undigested<br>Unfractionated |  | 5.00 | 12.88 | 15.63 | 14.38 |
| Serially<br>digested<br>fraction | Short | 3.50 | 16.00 | 1.63 | 4.38 |
|  | Long | 8.00 | 1.50 | 3.88 | 3.25 |
|  | Pellet | 16.13 | 11.63 | -5.38 | 3.38 |
|  | *Mean* | *9.21* | *9.71* | *0.04* | *3.67* |
| Histone<br>H2A<br>variant | H2A | 9.50 | 21.50 | -3.67 | 3.50 |
|  | H2A.B | 6.50 | 5.00 | 4.50 | 8.33 |
|  | macroH2A | 3.50 | -1.50 | 2.83 | 2.17 |
|  | H2B | 17.33 | 13.83 | -3.50 | 0.67 |

A notable difference between the viral and cellular DNA fragments was the presence of a di- and sometimes a tri-nucleosome peak in the digested cellular chromatin fractions, with deep valleys in between the peaks, as expected from stable nucleosomes, which were both replaced by a broad shoulder spanning from the mono-nucleosome peak in the viral DNA fragments, shown by the viral traces being above the valleys, and bellow the peaks, of the cellular ones in the short, long, and pelleted chromatin (Fig 4). In the undigested and unfractionated samples *(ND NF), the right side shoulder in the viral DNA was tighter than in the cellular DNA, as shown by the viral traces being consistently below the cellular ones on the right side from the peak (Fig 4), indicating that the broad shoulder of the viral DNA size distribution in the digested and immunoprecipitated fractions replaces the cellular di- and tri-nucleosome peaks (and intermediate valleys). This distribution is consistent with previous results showing a broader protection size for the viral mono- di- and tri-nucleosomes, which overlap each other (74, 75). The heterogeneity of the protected DNA is consistent with the concept that the viral nucleosomes are far more dynamic than the cellular ones, resulting in a broader distribution of the size of the fragments protected from limited (serial) digestion.

The differences in median or mode between the changes in immunoprecipitated over input viral and cellular DNA sizes were always at or below 12% (Table 5). Considering that HSV-1 DNA was co-immunoprecipitated with the histones in fragment sizes most compatible with those of the chromatinized cellular DNA in all fractions for all histones, therefore, we proceeded to quantitatively analyze the efficiency of assembly of the different histone H2A variants into H2A/H2B containing nucleosomes in the differentially accessible viral chromatin.

**Table 5.** Ratio of the median and mode length of the DNA fragments protected from MCN digestion and immunoprecipitated with anti-flag or anti-H2B antibodies over the total DNA fragments protected from MCN digestion by fraction (averages for the cell lines expressing any flag tagged histone) or histone (averages of all the fractions for each cell line), and of the relative changes in median and mode of the viral over the cellular DNA fragments. Ratios that differ by 10% or more are italicized, and by 12%, bolded.

| **Median** |  | **HSV-1**<br>(IP)/(No-IP) |  | **Cellular**<br>(IP)/(No-IP) |  | **Normalized**<br>[(HSV-1)/(Cell)] |  |
| --- | --- | --- | --- | --- | --- | --- | --- |
|  |  | Flag | H2B | Flag | H2B | Flag | H2B |
| Undigested<br>Unfractionated |  | 1.09 | 1.11 | 1.11 | 1.10 | 0.98 | 1.01 |
| Serially<br>digested<br>fraction | Short | 1.09 | 1.08 | 1.16 | 1.30 | 0.94 | 0.83 |
|  | Long | 1.07 | 1.06 | 1.19 | 1.18 | 0.90 | 0.90 |
|  | Pellet | 1.12 | 1.11 | 1.12 | 1.11 | 1.00 | 0.99 |
|  | *Mean* | 1.09 | 1.08 | 1.16 | 1.20 | 0.95 | 0.91 |
| Histone<br>H2A<br>variant | H2A | 1.10 | 1.11 | 1.14 | 1.23 | 0.97 | 0.90 |
|  | H2A.B | 1.08 | 1.08 | 1.15 | 1.17 | 0.94 | 0.92 |
|  | macroH2A | 1.08 | 1.04 | 1.17 | 1.19 | 0.93 | 0.88 |
|  | H2B | 1.10 | 1.10 | 1.16 | 1.18 | 0.95 | 0.93 |
| **Mode** |  |  |  |  |  |  |  |
| Undigested<br>Unfractionated |  | 1.03 | 1.09 | 1.09 | 1.09 | 0.94 | 1.00 |
| Serially<br>digested<br>fraction | Short | 1.02 | 1.09 | 1.01 | 1.03 | 1.01 | 1.07 |
|  | Long | 1.04 | 1.01 | 1.02 | 1.02 | 1.02 | 0.99 |
|  | Pellet | 1.09 | 1.06 | 0.97 | 1.02 | 1.12 | 1.04 |
|  | *Mean* | 1.05 | 1.05 | 1.00 | 1.02 | 1.05 | 1.03 |
| Histone<br>H2A<br>variant | H2A | 1.05 | 1.12 | 0.98 | 1.02 | 1.08 | 1.10 |
|  | H2A.B | 1.04 | 1.03 | 1.03 | 1.05 | 1.01 | 0.98 |
|  | macroH2A | 1.02 | 0.99 | 1.02 | 1.01 | 1.00 | 0.98 |
|  | H2B | 1.09 | 1.08 | 0.98 | 1.00 | 1.12 | 1.07 |

We first tested whether all cells supported HSV-1 DNA replication with similar efficacies and the viral DNA fractionated similarly in all (Fig 5 A, D, G, J). There were no differences in the total levels, or the fractionation, of HSV-1 DNA across the cells expressing the different flag tagged histones and thus the efficiency of co-immunoprecipitation in the different cell lines can be used to infer the fraction of nucleosomes containing the different H2A/H2B dimers.

**Figure 5.**
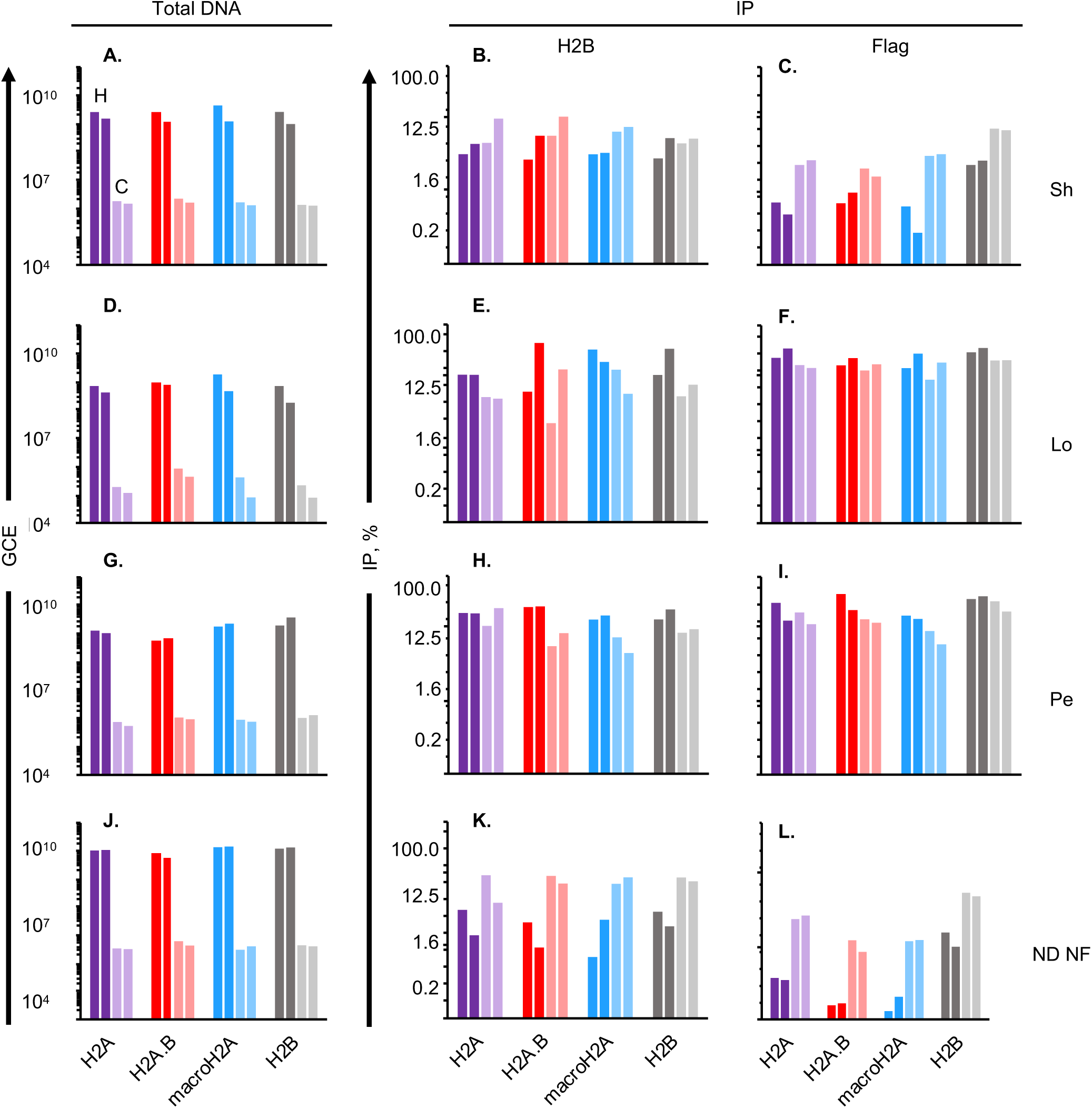
All tested histone H2A variants are assembled into HSV-1 chromatin regardless of chromatin dynamics. Chromatin of HeLa cells expressing each flag tagged histone and infected with HSV-1 for 7 hours was serially digested with MCN followed and fractionated was co-immunoprecipitated with anti-flag or anti-H2B antibodies. (A, D, G, J) Bar graphs presenting HSV-1 or cellular genome copy equivalents (GCE) for each cell line in each chromatin fraction from two independent biological infections. (B, C, E, F, H, I, K, L) Bar graphs presenting the log_2_ immunoprecipitation efficiency of HSV-1 or cellular DNA with anti-flag or anti-H2B antibodies from two independent biological infections. Sh, short accessible chromatin; Lo, long accessible chromatin; Pe, insoluble chromatin; ND NF, non-digested non-fractionated but sonicated chromatin. Dark shades, HSV-1; light shades, cellular; purple, cells expressing flag tagged H2A; red, cells expressing flag tagged H2A.B; blue, cells expressing flag tagged macroH2A; grey, cells expressing flag tagged H2B; H, HSV-1 DNA; C, cell DNA; GCE, genome copy equivalents.

HSV-1 DNA specifically co-immunoprecipitated with H2B and all tested flag tagged H2A variants in all fractions (Fig 5 B, C, E, F, H, I, K, L). HSV-1 DNA co-immunoprecipitated with flag antibodies far more efficiently in all chromatin fractions than with non-specific antibody; the average background HSV-1 genome copy number coimmunoprecipitated with non-specific mouse antibody for all chromatin fractions was between 6 and 13 % for each individual experiment, and 22 and 25 % for the unfractionated undigested chromatin. Non-specific background was subtracted before all analyses.

HSV-1 DNA specifically co-immunoprecipitated with H2B with similar efficiency in each fraction for each cell line, indicating no differences in assembly of HSV-1 chromatin (Fig 5 B, E, H, K). Interestingly, about 20-55 % of the HSV-1 or cellular DNA specifically co-immunoprecipitated with the flag or H2B antibodies in the different cell lines in the long soluble chromatin, and slightly less in the insoluble chromatin, whereas only about 1 to 10 % did in the short soluble chromatin, consistent with the concept that the latter contains the most dynamic viral nucleosomes. The overall lower efficiency of co-immunoprecipitation of the viral DNA with histones is thus a net effect of the enrichment of the viral DNA in the most dynamic chromatin fraction (2).

HSV-1 DNA specifically co-immunoprecipitated with H2B and all H2A variants except H2A.B with somewhat lower efficiency than with cellular DNA in the most accessible chromatin (Fig 6 B). By contrast, HSV-1 DNA specifically co-immunoprecipitated with H2A.B about 1.4-fold more efficiently than cellular DNA in this fraction, suggesting an enrichment in H2A.B containing nucleosomes specifically in the most dynamic viral chromatin. HSV-1 DNA specifically co-immunoprecipitated with H2A, H2A.B, or macroH2A with similar efficiency in all cell lines in the intermediately accessible chromatin (Fig 6 C, D), whereas it did so more efficiently than cellular DNA in the chromatin pellet (Fig 6 E, F), consistent with the concept that the least dynamic viral chromatin is highly enriched in the insoluble chromatin pellet.

**Figure 6.**
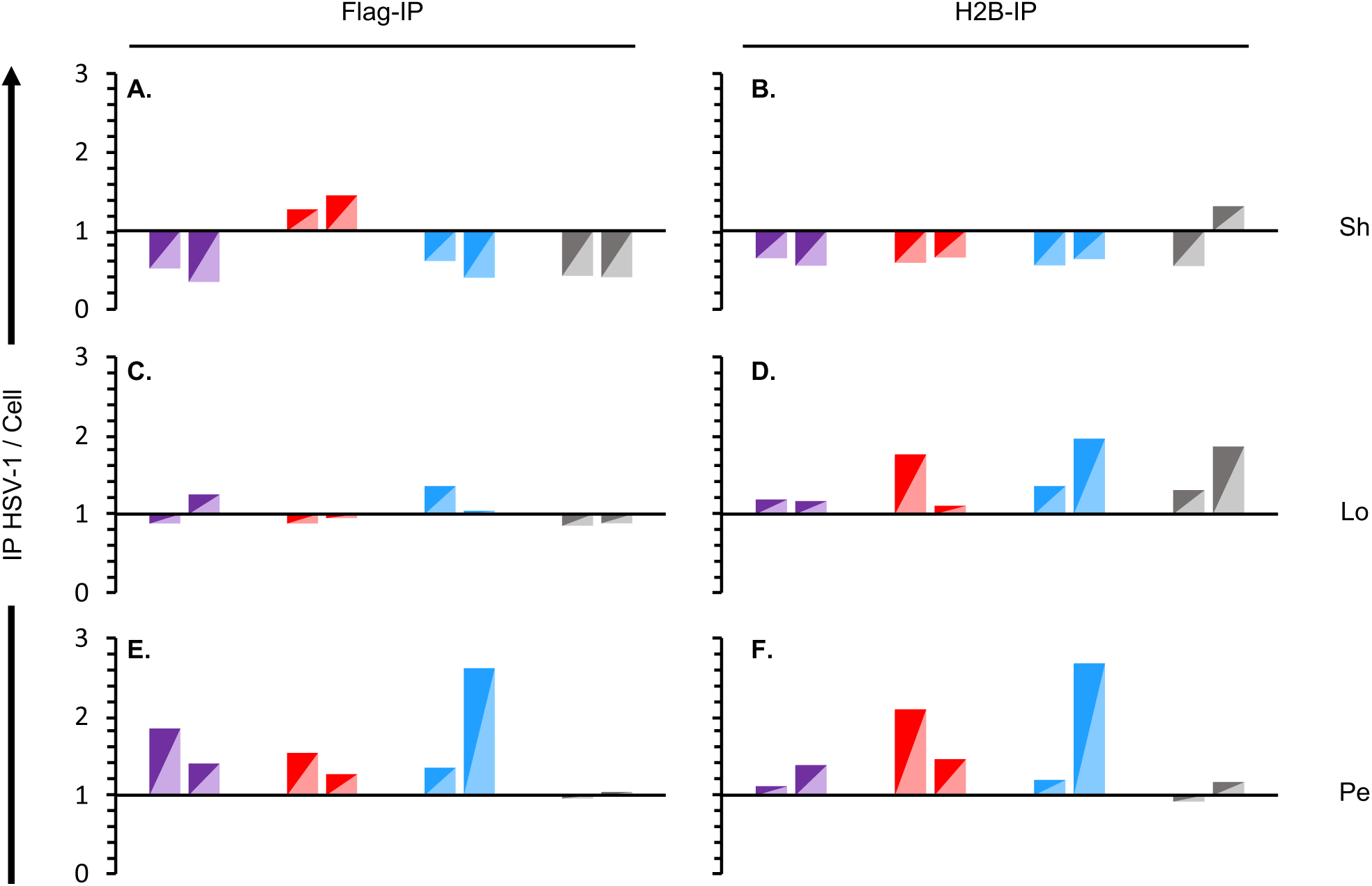
H2A.B co-immunoprecipitated with HSV-1 DNA with higher efficiency than with cellular DNA in the most accessible chromatin. Bar graphs presenting the ratio of HSV-1 to cellular DNA specifically co-immunoprecipitated with anti-flag (A, C, E, G) or anti-H2B (B, D, F, G) antibodies in different chromatin fractions from two independent biological experiments (biorepeats). Sh, short accessible chromatin; Lo, long accessible chromatin; Pe, insoluble chromatin; ND NF, non-digested non-fractionated but sonicated chromatin.

### Endogenous histone H2B and flag tagged histone H2A, H2A.B, macroH2A, or H2B are homogeneously distributed throughout the HSV-1 genome regardless of the dynamic state of the HSV-1 chromatin

The previous results indicate that different H2A variants are differentially incorporated into nucleosomes in the viral over the cellular chromatin of different accessibility. These differences could have resulted from a particular enrichment of H2A.B containing nucleosomes into more or less transcribed loci, or from a homogeneous enrichment of H2A.B containing nucleosomes across the genome in the most accessible viral chromatin. To differentiate between these possibilities, HSV-1 coverage plots were constructed from 50,000 HSV-1 aligned paired-end reads from chromatin co-immunoprecipitated with anti-flag or anti-H2B antibodies and normalized to the reads from the non-immunoprecipitated DNA in each fraction, to account for any potential sequence bias (Fig 7).

**Figure 7.**
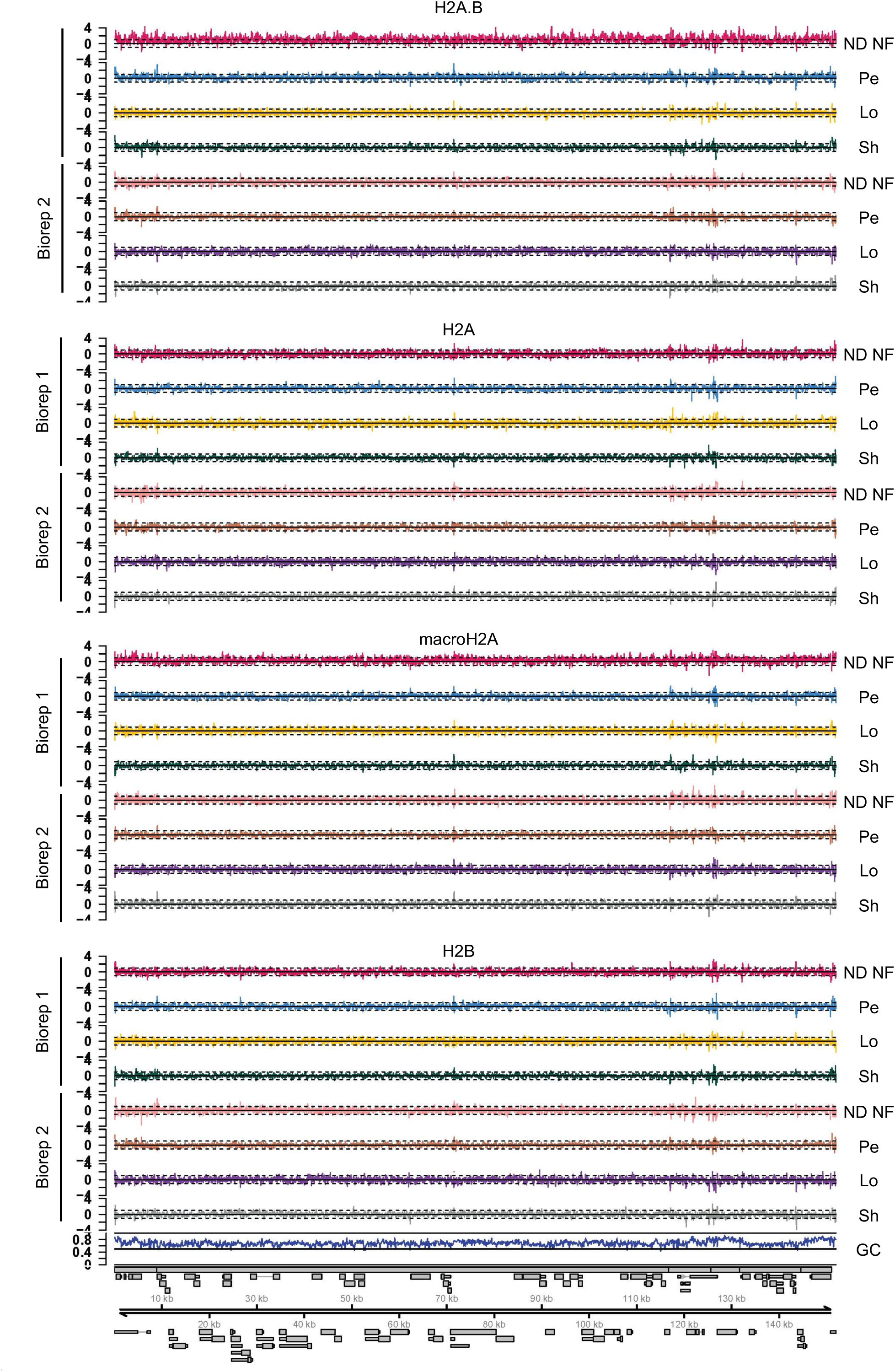
Endogenous histone H2B and flag tagged histone H2A, H2A.B, macroH2A, or H2B are homogeneously distributed throughout the HSV-1 genome regardless of the dynamic state of the HSV-1 chromatin. Line graphs presenting the log2 HSV-1 coverage (read depth) plots at single nucleotide resolution from 50,000 HSV-1-aligned paired-end reads normalized by the IP coverage [(Flag IP/No IP) / (H2B IP/No IP)].

Nucleosomes containing any H2A variant are assembled with H2B. To analyze the relative efficiency of assembly of the different H2A-H2B dimers into HSV-1 nucleosomes in the chromatin at different loci in the HSV-1 genome, we analyzed together the ChIP with anti-flag or anti-H2B antibodies. Anti-H2B antibody co-immunoprecipitates nucleosomes containing H2B in dimers with endogenous or the recombinant H2A variants, whereas the anti-flag antibody only co-immunoprecipitates nucleosomes containing tagged H2B or the specific H2A variant tagged in each cell line. We thus normalized the coverage of the distribution of each histone variant (Flag IP/No IP) to the coverage of H2B (H2B IP/No IP) (Fig 7). This normalization tests whether any H2A variant is preferentially enriched in the H2A/H2B dimers in the HSV-1 nucleosomes in any particular loci in any chromatin fraction, corrected by the efficiency of immunoprecipitation of the flag versus the H2B antibodies. If a particular variant were enriched in the H2A/H2B dimers in certain loci, then the ratio of that variant H2A to H2B will be higher than the average at those particular loci, and lower in the rest of the genome.

All histone H2A variant/H2B containing nucleosomes were similarly evenly distributed across the HSV-1 genome regardless of chromatin dynamics (Fig 7). The apparent enrichment across the genome for H2A.B in the unfractionated undigested chromatin in the first biological independent experiment is an artifact resulting from limiting number of reads in the input in that fraction. Downsampling to account for this difference brings this fraction to average about 1, at the cost of increased noise. Supporting figure 1 A-D presents the non-normalized coverage for the different H2A variant or H2B in each fraction for each cell line, indicating that there were no regions of the viral genome that had a lower efficiency of co-immunoprecipitation with any tested histone. H2A.B/H2B containing nucleosomes are thus preferentially incorporated into the most dynamic HSV-1 chromatin equally through the entire HSV-1 genome.

The ratio of HSV-1 DNA specifically co-immunoprecipitated with anti-flag or anti-H2B antibodies from cells expressing flag tagged H2B indicates the relative efficiency of incorporation of flag tagged H2B (Fig 8 A). For all other cells, this ratio indicates the relative efficiency of incorporation of each recombinant flag tagged H2A variant into the H2A/H2B dimers in the viral or cellular nucleosomes (Fig 8 A). Of note, these ratios are dependent on the percentage of the different H2A variants incorporated into the H2A/H2B dimers but also on the respective affinity of the anti-H2B and flag antibodies. Therefore, although these ratios can be used to comparatively analyze the different histone variants, they do not quantitate the actual percental composition of each H2A/H2B dimers, neither the percentage incorporation of tagged H2B.

**Figure 8.**
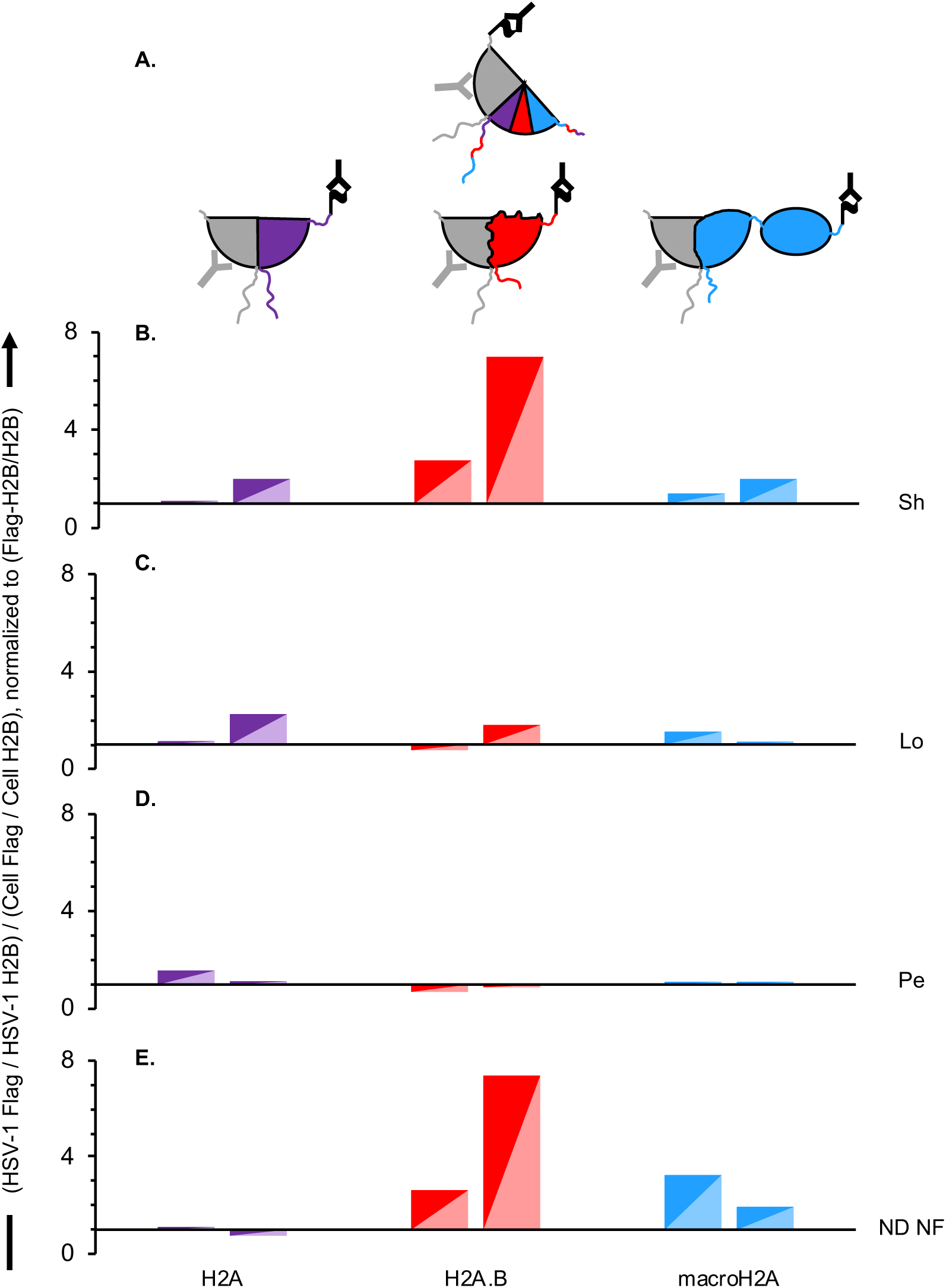
H2A.B is enriched in the most dynamic HSV-1 chromatin. (A) Cartoon depicting the different H2B-H2A dimers assembled in each cell line according to the ectopically expressed flag tagged histone. Purple, flag tagged H2A; red, flag tagged H2A.B; blue, flag tagged macroH2A; grey, flag tagged H2B or total H2B; black **Y**, anti-flag antibody; grey **Y**, anti-H2B antibody. (B, C, D, E) Bar graphs presenting the ratio of HSV-1 to cellular DNA co-immunoprecipitated with anti-flag or anti-H2B antibodies normalized to the ratio of HSV-1 or cellular DNA co-immunoprecipitated with flag tagged H2A variants corrected by the co-immunoprecipitation with flag tagged H2B or total H2B in different chromatin fractions from two independent biological infections. Sh, short accessible chromatin; Lo, long accessible chromatin; Pe, insoluble chromatin; ND NF, non-digested non-fractionated but sonicated chromatin.

We next analyzed the relative efficiency of incorporation of each flag tagged histone variant into nucleosomes in HSV-1 or cellular chromatin. To this end, we normalized the ratio of HSV-1 or cellular DNA co-immunoprecipitated with flag tagged histones to total H2B to the ratio of HSV-1 or cellular DNA co-immunoprecipitated with flag tagged H2B to total H2B (which corrects for the relative efficiency of immunoprecipitation by the anti-flag or anti-H2B antibodies). We then calculated the ratio of the normalized HSV-1 to cellular DNA co-immunoprecipitated with each H2A variant (Fig 8 B, C, D, E).

There was an enrichment of H2A.B/H2B containing nucleosomes in the total viral chromatin over the cellular one (Fig 8 E), which resulted from a relative enrichment of H2A/H2B containing nucleosomes in the viral chromatin specifically in the most accessible chromatin fraction (Fig 8 B). Canonical H2A, H2A.B, and macroH2A were in contrast similarly incorporated into the intermediate to least accessible viral or cellular chromatins (Fig 8 C, D).

### Histone H2A variants are differentially incorporated into the H2A/H2B dimers in the HSV-1 chromatin when HSV-1 transcription is modulated

We have previously proposed that the genomes in the most dynamic chromatin are the transcriptionally competent ones, although not necessarily they are entirely transcribed (2). The observed relative enrichment of H2A.B in the viral chromatin in this fraction through the entire genome at 7 hpi is consistent with this model, but all viral genes are transcribed at this time, precluding the analyses of non-transcribed loci.

We therefore tested next whether different levels of HSV-1 transcription in different loci affect the incorporation of H2A.B/H2B containing nucleosomes into HSV-1 chromatin. To this end, we modulated HSV-1 transcription with cycloheximide (CHX). CHX restricts HSV-1 transcription to the IE loci, resulting in a reduction of the fraction of HSV-1 genomes in the most accessible chromatin from about a third to about 2 % (2). As the previous experiments had shown that the total incorporation of the different histone variants in the unfractionated viral genomes directly reflected the enrichment of H2A.B/H2B containing nucleosomes in the most accessible viral chromatin (Fig 8), and the percentage of HSV-1 genomes in the most accessible chromatin decreases to only about 2 % in the presence of CHX (2), we pursued analyses of total unfractionated chromatin.

Chromatin of cells expressing each flag tagged histone and infected with HSV-1 for 7 hours, treated or not with CHX, was specifically co-immunoprecipitated with anti-flag or -H2B antibodies and sequenced. All cells supported HSV-1 DNA replication with similar efficacies, both in the absence, as before (Fig 5), or presence of CHX (Fig 9 A, B). We thus proceeded to evaluate co-immunoprecipitation efficiency. We lost one sample to an unidentified sequencing issue that resulted in an abnormally high number of total reads together with an abnormally low percentage of viral reads in the macroH2A expressing cell line immunoprecipitated with H2B antibody in the CHX treatment (Biorepeat 1); this sample was not analyzed.

**Figure 9.**
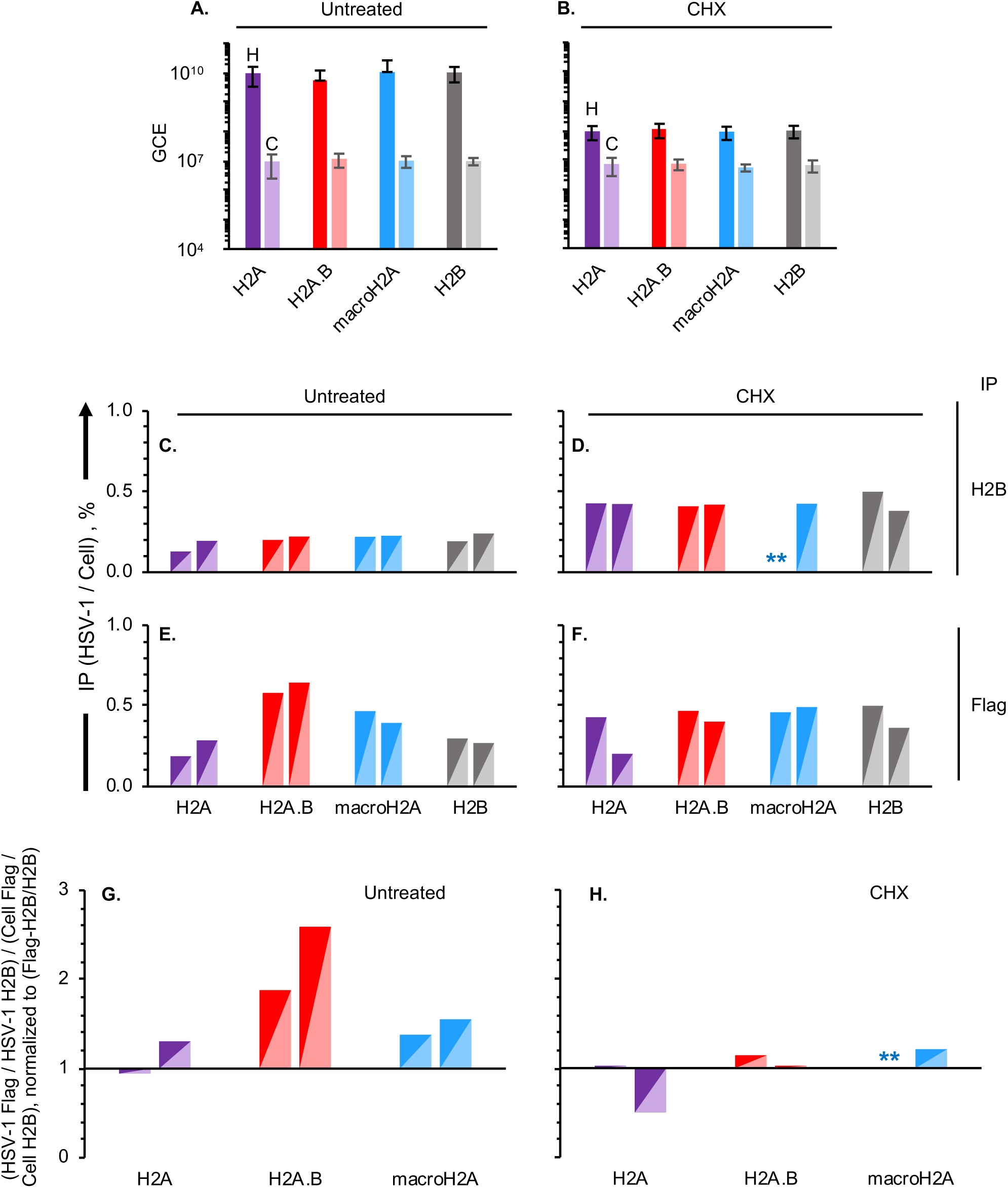
H2A.B is not enriched in the HSV-1 chromatin when transcription is restricted to the IE loci. Chromatin of cells expressing flag tagged H2A, H2A.B, macroH2A, or H2B, HSV-1 infected, pre-treated with cycloheximide for one hour or not harvested at 7 hours after infection in the presence of CHX or not was co-immunoprecipitated with anti-flag or anti-H2B antibodies. (A, B) Bar graphs presenting HSV-1 or cellular genome copy equivalents (GCE) in each cell line from four independent untreated (A) or three cycloheximide-treated (B) infections. (C, D, E, F) (C-F) Bar graphs presenting the ratio of HSV-1 to cellular sequenced DNA co-immunoprecipitated with anti-H2B (C, D) or anti-flag (E, F) antibodies from two independent untreated (C, E) or cycloheximide-treated (D, F) infections. (G, H) Bar graphs presenting the ratio of HSV-1 to cellular reads DNA co-immunoprecipitated with anti-flag or anti-H2B antibodies normalized to the ratio of HSV-1 or cellular sequenced DNA co-immunoprecipitated with flag tagged H2A variants corrected by the co-immunoprecipitation with flag tagged H2B or total H2B from two independent untreated (G) or cycloheximide-treated (H) infections. Dark shades, HSV-1; light shades, cellular; purple, cells expressing flag tagged H2A; red, cells expressing flag tagged H2A.B; blue, cells expressing flag tagged macroH2A; grey, cells expressing flag tagged H2B; H, HSV-1 DNA; C, cell DNA; GCE, genome copy equivalents; CHX, cycloheximide.

HSV-1 DNA co-immunoprecipitated with H2B 13 to 24 % as efficiently as cellular DNA when viral transcription was unrestricted and the genomes are in the most dynamic chromatin (Fig 9 C), very much as in the previous experiments (Fig 5). Under conditions when HSV-1 transcription is limited to the IE loci and the viral chromatin is less dynamic (2), however, HSV-1 DNA co-immunoprecipitated with H2B more efficiently, at about 38 to 50 % of that of cellular DNA (Fig 9 D). Co-immunoprecipitation with flag tagged H2A variants was differentially efficient at about 19 to 29 % of that of cellular DNA for H2A, 39 to 47 % for macroH2A, or 58 to 64 % for H2A.B in untreated infections (Fig 9 E), again consistent with the previous experiments (Fig 5, 6). Co-immunoprecipitation of HSV-1 DNA with flag tagged H2A variants increased by 34 to 50 % for H2A and H2B and about 10 % for macroH2A, but decreased for H2A.B by about 30 % (Fig 9 D, F), indicating that restriction of transcription and enrichment of the HSV-1 DNA in the least accessible chromatin (2) result in a differential effect on its assembly into nucleosomes with the different histone H2A variants.

To analyze the relative efficiency of genome-wide incorporation of each flag tagged histone H2A variant into H2A/H2B containing nucleosomes in HSV-1 or cellular chromatin, we normalized the ratio of HSV-1 or cellular DNA co-immunoprecipitated with flag tagged histones to total H2B to that of HSV-1 or cellular DNA co-immunoprecipitated with flag tagged H2B to total H2B as before. We then calculated the ratio of the normalized HSV-1 to cellular DNA co-immunoprecipitated with each H2A variant (Fig 9 G, H), as before (Fig 8). H2A.B was again preferentially incorporated into the H2A/H2B nucleosomes in viral chromatin over H2A or macroH2A when all viral genes are transcribed (Fig 9 G), as before (Fig 8), but not when transcription was restricted and HSV-1 DNA becomes less accessible (Fig 9 H).

### Endogenous histone H2B and flag tagged histone H2A, H2A.B, macroH2A, or H2B are homogeneously distributed throughout the HSV-1 genome even when transcription is restricted to the IE loci

HSV-1 DNA was no longer co-immunoprecipitated with higher efficiency with H2A.B in the most dynamic chromatin when transcription was restricted to the IE loci, but this lack of global enrichment could result from an overall depletion of H2A.B/H2B containing nucleosomes through the genome together with a particular enrichment in certain loci, like the transcribed IE ones. To test the distribution of the different histone H2A variant/H2B containing nucleosomes across the HSV-1 genome, HSV-1 coverage plots were constructed from 50,000 HSV-1 aligned paired-end reads from chromatin co-immunoprecipitated with anti-flag or anti-H2B antibodies, normalized to the reads from the input DNA before immunoprecipitation to account for any sequencing bias (Fig 10).

**Figure 10.**
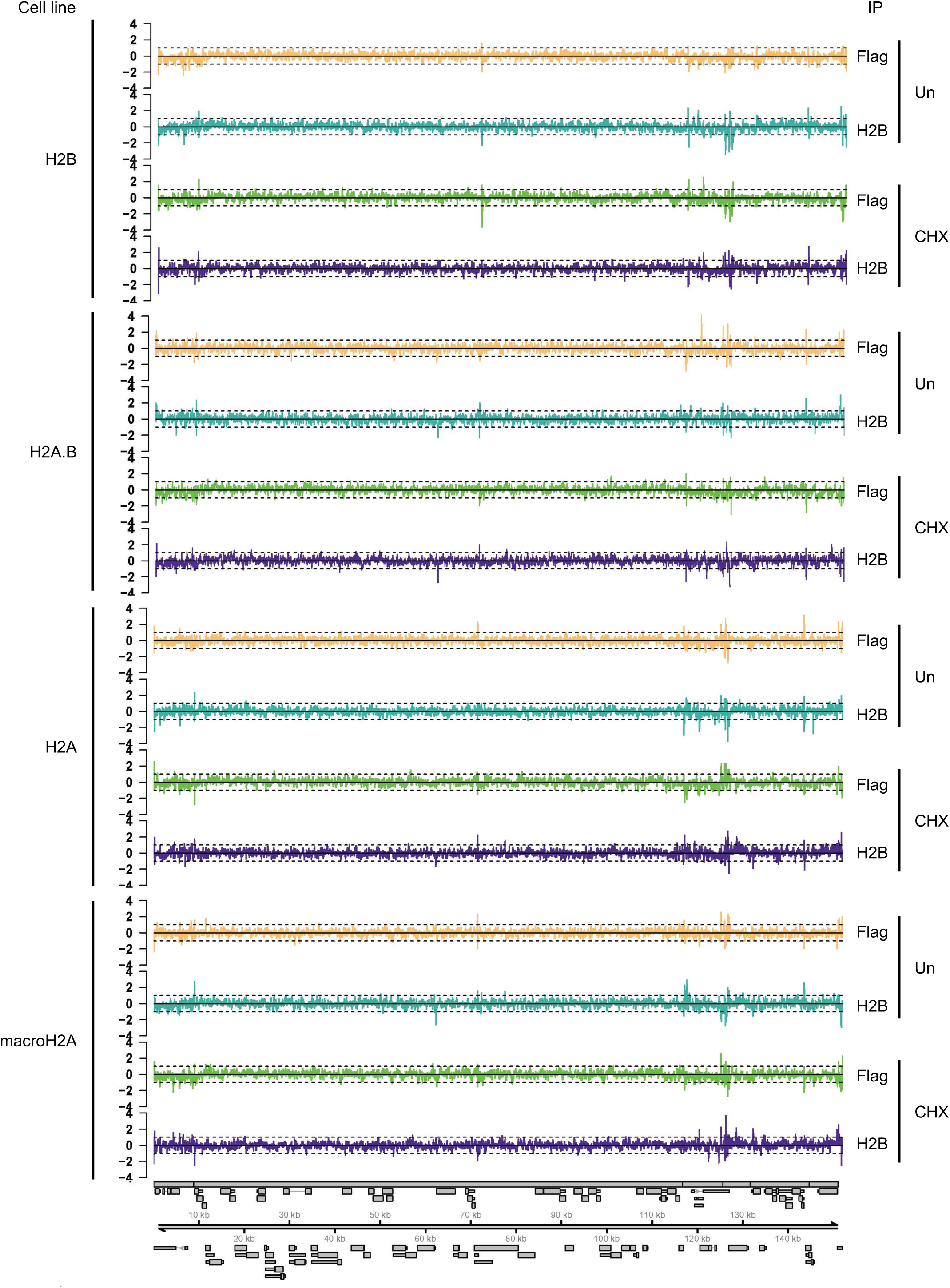
No H2A variant is enriched or depleted from the IE loci when transcription is restricted to the IE genes only. Line graphs presenting the log2 HSV-1 coverage plots from 50,000 HSV-1-aligned paired-end reads normalized by the IP coverage [(Flag IP/No IP) / (H2B IP/No IP)]; Un, untreated infection; CHX, cycloheximide-treated infection; Gray rectangles, cartoon representation of the HSV-1 genome.

Restriction of HSV-1 transcription to the IE loci had no obvious effect on the distribution of H2B or any H2A variants across the HSV-1 genome. After normalizing the coverage of each variant to that of H2B as before, to evaluate the distribution of H2A/H2B containing nucleosomes (Flag IP/No IP normalized to H2B IP/No IP), all H2A variant/H2B containing nucleosomes were similarly distributed across the HSV-1 genome regardless of whether transcription was restricted to the IE loci or ongoing throughout the HSV-1 genome (Fig S2). The also even distribution of the FlagH2B normalized to total H2B indicates that the incorporation of flag tagged histones is not biased toward, or away from, different loci of the genome even when transcription is restricted to the IE loci. The also even distribution of the non-normalized co-immunoprecipitation with any flag tagged H2A or H2B or endogenous H2B indicates that no loci in the genome is preferentially chromatinized or non-chromatinized.

We conclude that transcriptionally active HSV-1 lytic chromatin is preferentially assembled into H2A.B/H2B containing nucleosomes whereas the transcriptionally repressed one is not (Fig 11).

**Figure 11.**
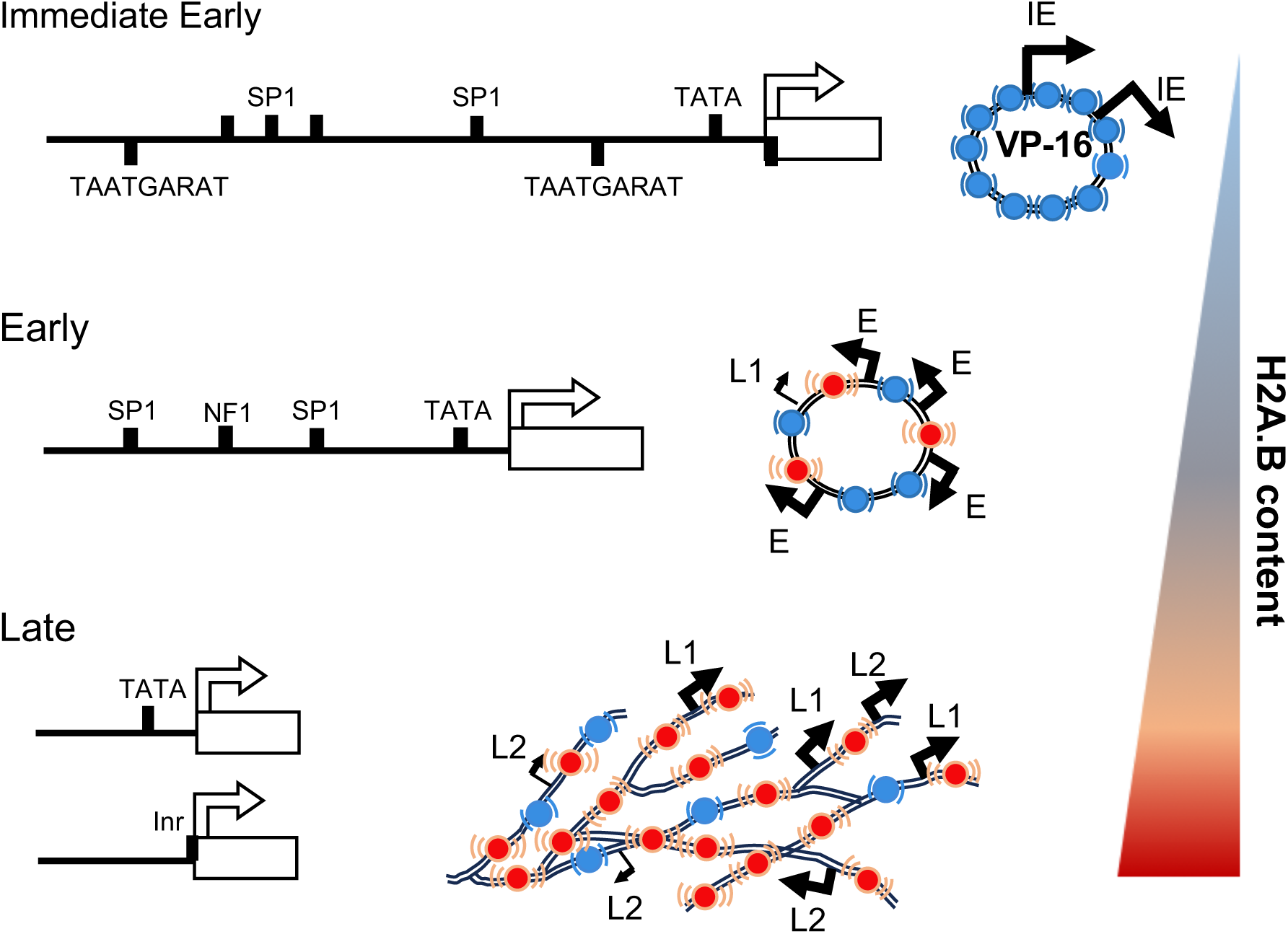
H2A.B is preferentially incorporated into highly dynamic and transcriptionally competent HSV-1 chromatin but not into stable and transcriptionally repressed HSV-1 chromatin. Cartoon depicting the enrichment in HSV-1 chromatin assembled with H2A.B containing nucleosomes as infection progresses and the transcriptional competence of the genomes for transcription of IE, E and L genes when the chromatin is differentially enriched in H2A.B.

## Discussion

Here we show that the HSV-1 lytic chromatin is enriched in histone variant H2A.B, which is associated with highly dynamic and transcriptionally competent chromatin (95). H2A.B is a minor inducible H2A variant, except in testis, which among somatic tissues is expressed to somewhat higher levels in skin less so in sensory or dorsal root ganglia (50). It is enriched in the nucleoli and in highly transcribed genes (60, 93). In cultured human primary cells, it is expressed to higher levels in those that support HSV-1 gene expression (52), and its mRNA levels increase during infection (53). H2A.B dynamics decreased in infected cells consistently with a larger fraction spending more time assembled into chromatin with the novel binding sites provided by HSV-1 DNA.

MCN digestion in the serial protocol is performed on native chromatin. The lower efficiency of HSV-1 DNA co-immunoprecipitation in the most accessible chromatin could indicate that there are fewer nucleosomes in the viral DNA in this fraction or that the interactions are more dynamic. The former would be expected to result in lower recovery of viral than cellular DNA and a larger frequency of smaller DNA fragments. The latter would be reflected in a quantitative recovery of the viral DNA and an approximately normal, although broad, fragment size distribution. Viral DNA was quantitatively recovered, and even more so in the cells expressing flag-tagged H2A.B, indicating similar level of protection to mild nuclease digestion of the viral and cellular DNA. The size distribution of the protected viral and cellular fragments had similar short left and long right tails. There were the same very low frequencies of reads shorter than 100 bp for the viral as for the cellular DNA. The mean and mode of the protected viral DNA in the most accessible viral chromatin fraction were also basically identical to those of the cellular DNA, even a bit longer, indicating that both DNA were protected in nucleoprotein complexes involving approximately the same length of DNA, which would not be expected if a large fraction of the viral DNA was non-nucleosomal. The HSV-1 DNA co-immunoprecipitated by any histone had a similar increase in median and mode as the cellular DNA, and not much difference in the fragment size distribution, other than the broad right shoulder replacing the peaks and valleys. A model proposing large stretches of non-nucleosome DNA would have predicted a much larger increase in the mean, and a much tighter distribution, with sharper mode, in the co-immunoprecipitated than the input viral DNA. The mechanism most consistent with the observed results is therefore HSV-1 DNA being protected by its assembly into most dynamic nucleosomes, which due to their innate instability disassemble after digestion and before cross linking, resulting in the lower percentage of co-immunoprecipitation.

Classical protection assays of HSV-1 lytic chromatin fail to produce the typical “nucleosome ladder” (74). The nucleosome ladder results from the difference in histone dynamics. Linker histone H1 has a much shorter residency time than the core nucleosome (89, 96). Therefore, linker DNA is exposed and cleaved far more frequently than nucleosome DNA. Tagmentation to completion of HSV-1 DNA does not produce the typical decreasing sigmoidal distribution of tagmented fragments depleted in multiples of mono-to poly-nucleosome size either. Instead, the pattern is more consistent with naked DNA or dynamic chromatin in which both linker and nucleosome DNA are similarly accessible, producing an enrichment DNA fragments of subnucleosomal size followed by a steady decline in the frequency of tagmentation of longer DNA fragments (25, 85, 97). If the dynamics of linker and core histone were to become more similar, then the protection patterns to nuclease digestion or tagmentation would change to heterogeneously sized fragments resolving as smears in agarose gels. H2A.B is the most dynamic core histone, having a residency time closer to that of linker histone H1 than all other core histones (32, 98) (Fig 1). The DNA in H2A.B containing nucleosome is wrapped around less stably than in H2A containing ones, making it more accessible (61, 72, 88). Nucleosomes assembled with H2A.B are consequently protected in broader bands (72, 73, 88), plus heterogeneously sized fragments (88).

The serial digestion and fractionation of HSV-1 lytic chromatin presented here show that the cellular DNA fragments had the expected distribution as mono-to poly-nucleosome sizes with marked valleys in between, and a very sharp mode of 167 bp in the most accessible chromatin, highly enriched in mono-nucleosomes. Consistently with previously published results (74, 85, 97), the far more dynamic viral chromatin resulted in DNA fragments with a broader distribution and without clear valleys in between the different poly-nucleosome sizes. Nonetheless, the changes in both the mode and the mean of the HSV-1 DNA fragments protected from mild (serial) MCN digestion and co-immunoprecipitated with any ectopically expressed H2A variant tested or ectopically expressed or native H2B were all but three in between 12 %, and all but seven in between 10 %, of the changes in the lengths of the cellular DNA fragments (Table 5).

The observed enrichment in H2A.B in the viral chromatin is consistent with the patterns of protection of the HSV-1 DNA to nuclease digestion or tagmentation, and the distribution of the protected fragments is inconsistent with long stretches of non-chromatinized HSV-1 DNA.

Selected HSV-1 loci in lytically infected cells co-immunoprecipitates with all core histones (81), including H3 bearing a variety of post translational modifications (PTM) (2, 4, 9–11, 14, 21–24, 79, 81, 82, 85, 99–106). The entire HSV-1 genome co-immunoprecipitates with H3 (79, 85), H2A, and H2B (Figs 4-10), although the interactions between histones and HSV-1 DNA have been described as sparser, weaker, or more dynamic than their interactions with cellular DNA (2, 10, 11, 74, 75, 81, 83, 107, 108). Total or post-translationally modified histone H3 is widely and evenly spread throughout the HSV-1 genome, but the efficiency of co-immunoprecipitation changes as the infection progresses and HSV-1 chromatin dynamics increase (2, 79, 85). Likewise, here we show that the efficiency of HSV-1 DNA co-immunoprecipitation with H2B and H2A also increases when transcription is restricted to the IE loci, and the viral chromatin is less dynamic (2). Considered together, these results indicate that the dynamics of the viral chromatin increase across the genome as transcription increases across the genome.

Canonical H3.1 and variant H3.3, which assemble the least and most dynamic nucleosomes respectively, are unstably assembled in HSV-1 chromatin (2, 66, 109), but H3.3 is assembled in HSV-1 chromatin immediately upon nuclear entry whereas H3.1 does so only after the onset of HSV-1 DNA replication (109). The dynamics of both H3.1 and H3.3 are enhanced in HSV-1 infected cells, but the dynamics of only H3.1 are enhanced to an even greater extent when HSV-1 DNA replication is inhibited (91). HSV-1 RNA levels are lower when the assembly of H3.3 containing nucleosomes is inhibited (109). It is thus likely that the most dynamic nucleosomes containing H3.3 are less restrictive to HSV-1 transcription than the least dynamic ones containing H3.1. Likewise, the most dynamic nucleosomes containing H2A.B may be the least restrictive to HSV-1 transcription.

Canonical or any variant H2A all dimerize with H2B, whereas H2B dimerizes with either H2A or one of its many variants. Therefore, the dynamics of H2B are not necessarily expected to follow the same pattern as those of any single H2A but would rather reflect the average of the dynamics of all H2A variants. Nonetheless, the fast exchange rate of H2B decreased in cells infected with HSV-1 (90), as did that of H2A.B (Fig 1 D), but not that of canonical H2A (Fig 1 D, (31)) or other H2A variants (Fig 1 D). H2A, macroH2A, and H2A.X had faster fast exchange rates in HSV-1 infected cells whereas H2A.B had a slower one (Fig 1 D, E), most closely mimicking H2B dynamics again (90). As H2A/H2B dimers typically exchange as a unit, these similar dynamics are consistent with H2A.B being enriched in the H2A/H2B dimers in the nucleosomes in the viral chromatin and thus being the variant that influences the most the dynamics of H2B in the viral chromatin.

The lack of enrichment in H2A.B in the viral chromatin when transcription was restricted by CHX most likely directly reflects the large decrease of the fraction of HSV-1 genomes in the most accessible chromatin fraction (2), which is enriched in H2A.B (Fig 8). A caveat of these population analyses is that they cannot differentially evaluate the composition of the chromatin in a minority of genomes from which the IE genes may be transcribed from the vast majority from which they may not. We therefore cannot exclude that the chromatin of the IE loci in a subset of genomes may be enriched in H2A.B and that these may be the genomes from which the IE genes are transcribed.

The dsDNA genomes of HSV-1 are not chromatinized in the virion (86, 110, 111) but become assembled in highly dynamic chromatin during lytic infection (2, 74). After HSV-1 DNA replication, viral DNA accounts for about 20 % of the total nuclear DNA (2). Core nucleosomes assemble with 147 bp of dsDNA and are linked by up to 80 bp of dsDNA. The cellular genome has 3 billion bps, providing approximately 1.3×10^7^ nucleosome assembly sites. HSV-1 genomes are about 152 kbp long, providing approximately 675 new nucleosome assembly sites. At 7 hpi there are approximately 1,300 HSV-1 genomes in the nucleus of an infected cell, which would provide about 877,500 new nucleosome assembly sites, an increase of about 7%. The dynamics of all histones could have thus been expected to somewhat decrease during HSV-1 infection, as the number of histones is constant, but the number of binding sites increases. However, only H2A.B dynamics decreased. The dynamics of macroH2A, H2A , and H2A.X likely increase during HSV-1 infection as they are disassembled from the most numerous stable nucleosomes with cellular DNA to be reassembled in unstable nucleosomes with the replicating HSV-1 genomes. In contrast, the dynamics of H2A.B decrease, as H2A.B is only a minor variant and thus the H2A.B disassembled from highly dynamic cellular nucleosomes is promptly reassembled in, still unstable, nucleosomes with the newly present HSV-1 DNA.

In conclusion, transcriptionally competent lytic HSV-1 chromatin is enriched in nucleosomes containing histone variant H2A.B, which is associated with highly dynamic and transcriptionally active chromatin. The different H2A variants are assembled into HSV-1 chromatin evenly through the entire genome, rather at the loci of individual HSV-1 genes. The relative enrichment in the dynamic variant H2A.B would thus promote global HSV-1 transcriptional competency, but not regulate the transcription level of individual genes.

## Materials and Methods

### Cells

African green monkey kidney Vero cells (CCL-81, ATCC, distributed by Cedarlane Laboratories Ltd., ON, CA, or CRL-1587 obtained directly from BEI), were maintained in Dulbecco’s modified minimum Eagle’s medium (DMEM, 11885, Invitrogen, Burlington, ON, CA, or ThermoFisher Scientific [Gibco], Waltham, MA, USA) supplemented with 5 % fetal bovine serum (FBS, A15-70, PAA Laboratories Inc., Etobicoke, ON, CA) at 37°C in 5 % CO2. HeLa cells (ATCC) were maintained in Dulbecco’s modified minimum Eagle medium (DMEM) (Gibco) supplemented with 5 % fetal bovine serum ([FBS], Avantor Seradigm, VWR). Human embryonic kidney 293 cells expressing the SV40 T-antigen ([HEK-293T], CRL11268, ATCC) were maintained in DMEM supplemented with 5 % FBS.

### Viruses

Human herpesvirus 1 (HSV-1), strain KOS, passage 9, obtained from the late Dr. P. Schaffer, University of Pennsylvania, Philadelphia, PA, USA is described (112–114). A separate stock from ATCC (VR-1493) was used at passage 3 for the chromatin immunoprecipitation (ChIP) studies. Viral stocks were prepared and titres were determined as described (89–91).

### Drugs

Cycloheximide (CHX) 5 mg/ml stock solution was prepared in DMEM and diluted to 50 μg/ml in DMEM, DMEM containing 5 % FBS or PBS before use. Cells were pre-treated with 50 μg/ml CHX in DMEM containing 5 % FBS for 1 h prior to infection. CHX was maintained in the inocula, PBS or DMEM washes and DMEM containing 5 % FBS until harvest.

### Plasmids

pEGFP-H2B expression plasmid is described (90). To construct the pEGFP-H2A expression plasmid, the DNA sequence encoding H2A was PCR amplified (Table 6) from cDNA clone PX00928G15 obtained from the Riken Mouse cDNA library (mouse [AK028026] and human [AY131983] histone H2A encode for identical proteins). Amplified DNA encoding the sequence for H2A was ligated in frame with the BglII and SalI restriction digest sites of pEGFP-C1 (Clontech). To construct the pEGFP-H2A.X expression plasmid, the DNA sequence encoding H2A.X (H2AFX) was PCR amplified (Table 6) from a plasmid generously provided by Dr. Anette U. Duensing (Department of Pathology, University of Pittsburgh, Pennsylvania, US) (115). Amplified DNA encoding the sequence for H2A.X was ligated in frame with pEGFP-C1 (Clontech) at the BglII and SalI restriction sites.

**Table 6.**
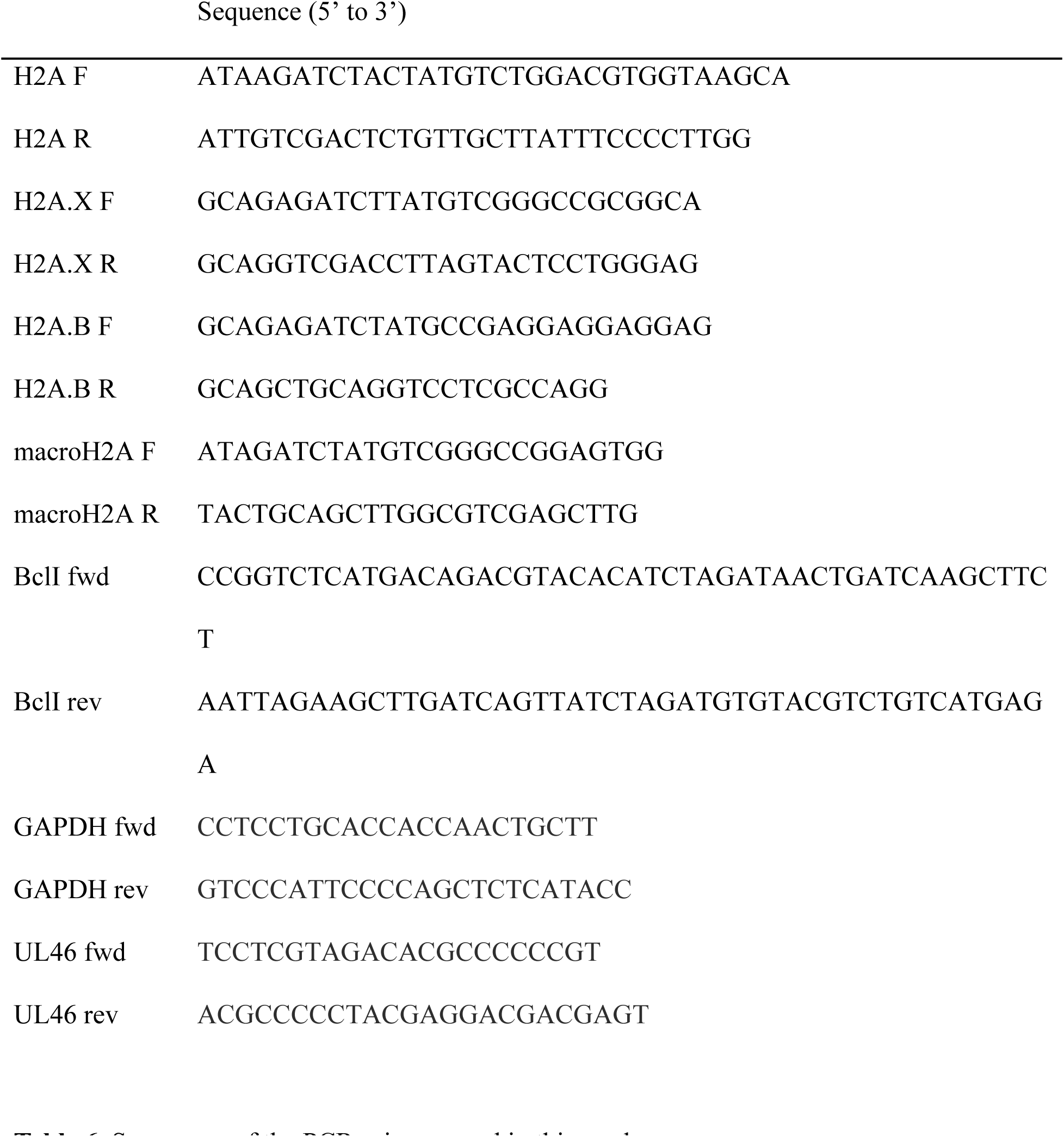
Sequences of the PCR primers used in this work.

To construct the pEGFP-H2A.B (H2A.B variant histone 2, H2AB2) expression plasmid, the DNA sequence encoding H2A.B was PCR amplified (Table 6) from a plasmid generously provided by Dr. Vasily Ogryzko (Institut Gustave Roussy, Villejuif, France) (60, 93). The H2A.B gene (H2A.B variant histone 2, H2AB2) was ligated in frame with the BglII and PstI restriction digest sites of pEGFP-C1 (Clontech). To construct the pEGFP-macroH2A1.2 expression plasmid, the DNA sequence encoding human macroH2A1.2 (H2AFY) was PCR amplified from a plasmid generously provided by Dr. Vasily Ogryzko (Institut Gustave Roussy, Villejuif, France) (60, 93). The macroH2A gene was ligated in frame with the BglII and PstI restriction sites of pEGFP-C1 (Clontech).

To construct the pH2A-3Xflag, pH2B-3Xflag and pH2A.B-3Xflag expression plasmids, the DNA sequence encoding EGFP was removed from pEGFP-H2A, pEGFP-H2B, and pEGFP-H2A.B by AgeI and BspEI restriction digestion followed by ligation of the compatible ends to produce plasmids pH2A, pH2B and pH2A.B. To insert the 3Xflag-tag sequence into the pH2A, pH2B and pH2A.B plasmids, gBlock DNA containing a sequence expressing DYKDHDG•DYKDHDI•DYKDDDDK (3Xflag-tag) (Integrated DNA Technologies [IDT]) (Table 7), followed by a stop codon and flanked by SalI and BamHI restriction digest sites, was ligated in frame with pH2A, pH2B, or pH2A.B at SalI and BamHI sites.

**Table 7.**
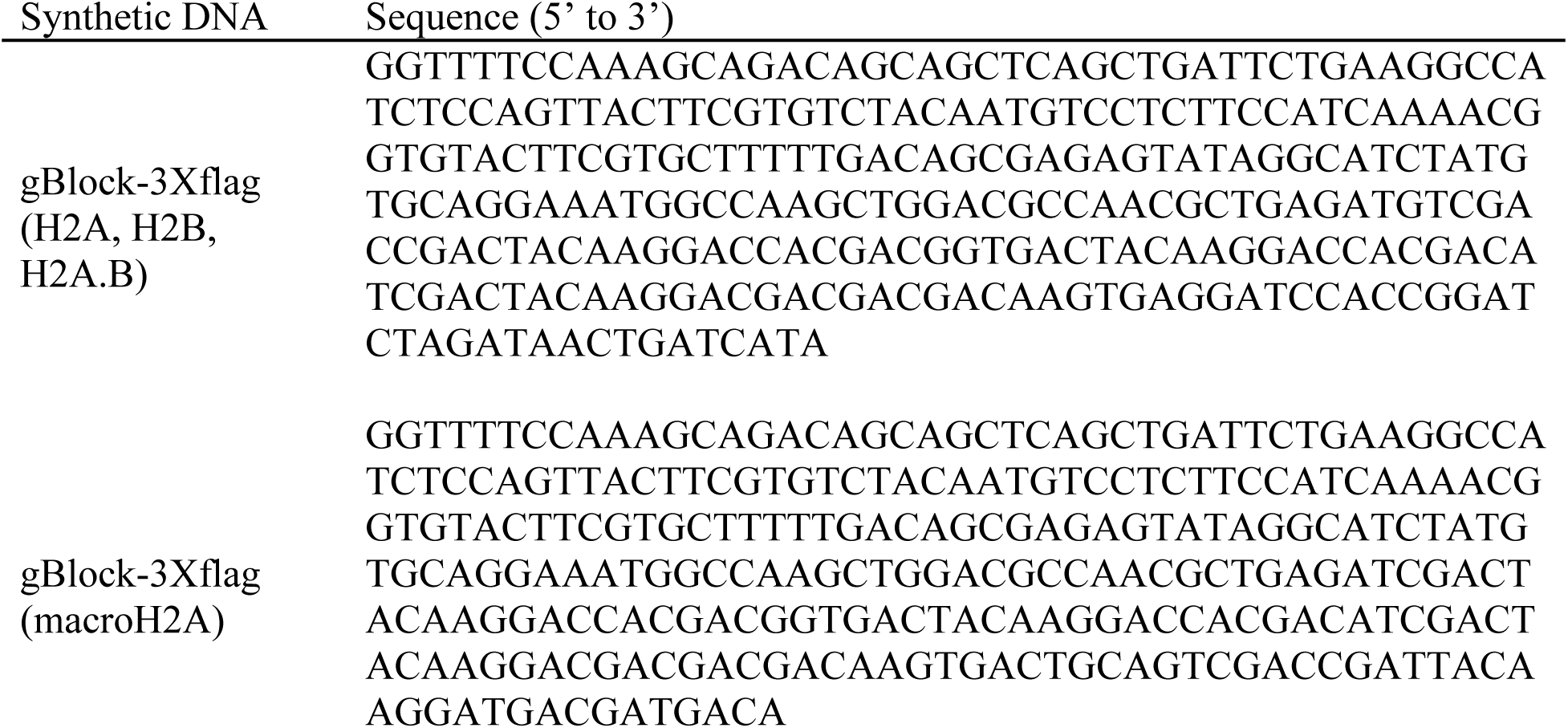
Sinthetic DNA used in the cloning of the different histone variants.

To construct the pmacroH2A-3Xflag expression plasmid, the DNA sequence encoding EGFP was removed from pEGFP-C1 by AgeI and BspEI restriction digest followed by ligation of the compatible ends to produce plasmid p-C1. macroH2A was transferred from pEGFP-macroH2A into p-C1 by ligating in frame at the BglII and PstI sites to produce plasmid pmacroH2A-C1. To insert the 3Xflag-tag sequence into pmacroH2A-C1, the gBlock (3Xflag-tag) DNA sequence (IDT) (Table 7), and stop codon, flanked by BlpI and SalI restriction digest sites, was ligated in frame with pmacroH2A-C1 at the BlpI and SalI restriction sites.

### Cloning of flag and EGFP tagged histones into lentivirus transfer vectors

The pLJM1-BclI lentivirus transfer vector was constructed by removing EGFP from pLJM1-EGFP (Addgene) by AgeI and EcoRI restriction digest, and then adding in frame complementary oligonucleotides BclI fwd and BclI rev (IDT) (Table 6), which contain the BclI cleavage site and overhangs compatible with AgeI and EcoRI sites. Plasmids pEGFP-H2A, pEGFP-H2B, pEGFP-H2A.B, pEGFP-macroH2A, pH2A-3Xflag, pH2B-3Xflag, pH2A.B-3Xflag, pmacroH2A-3Xflag and pLJM1-BclI were grown in dam^-^/dcm^-^ competent E.coli (NEB). EGFP-H2A, EGFP-H2B, EGFP-H2A.B, H2A-3Xflag, H2B-3Xflag, and H2A.B-3Xflag were extracted from pEGFP-H2A, pEGFP-H2B, pEGFP-H2A.B, pH2A-3Xflag, pH2B-3Xflag, and pH2A.B-3Xflag, respectively, and ligated in frame at NheI and BclI sites in pLJM1-BclI to produce pLJM1-EGFP-H2A, pLJM1-EGFP-H2B, pLJM1-EGFP-H2A.B, pLJM1-H2A-3Xflag, pLJM1-H2B-3Xflag, and pLJM1-H2A.B-3Xflag lentivirus transfer vectors. EGFP-macroH2A and macroH2A-3Xflag were extracted from pEGFP-macroH2A and pmacroH2A-3Xflag into pLJM1-BclI, respectively, and ligated in frame with AfeI and BclI sites of pLJM1-BclI to produce pLJM1-EGFP-macroH2A and pLJM1-macroH2A-3Xflag lentivirus transfer vectors, respectively.

### Lentiviral production

For each transfer vector, 5 x 10^5^ HEK-293T cells were seeded into 100 mm dishes. About 18 h later, 6 µg of 4:2:1:1 transfer:Gag-Pol (pMDLg-pRRE, Addgene):Rev (pRSV-Rev, Addgene):envelope (pMD2.G, Addgene) plasmids were mixed in 500 µl of Opti-MEM I reduced serum medium (Gibco) and incubated with 18 µl of TransIT-LT1 Reagent (Mirus) for 30 min at room temperature. Mixes were added dropwise to each dish spreading them throughout the entire surface and incubated at 37 °C for about 16 h, when media were replaced with 5 ml of 5 % FBS-DMEM. Forty-eight and seventy-two hours after transfection, 5 ml of supernatant was collected, and stored at 4 °C, and 5 ml of fresh 5 % FBS-DMEM was added. Harvested medium was cleared 15 min at 4,000 x g at 4 °C. Three milliliters of Lenti-X concentrator (Takarabio) and 9 ml of clarified supernatant were gently mixed by inversion, incubated at 4 °C for 30 min, pelleted at 1,500 x g for 45 min at 4 °C, and resuspended in 350 µl of DMEM, aliquoted, frozen in ethanol/dry ice, and stored at – 80 °C.

### Lentiviral titration

We used the matching EGFP tagged histones as a proxy to estimate the titer of the lentivirus encoding flag tagged histones. Thirty thousand HeLa cells were seeded into each well of a 48 well plate. Five tenfold viral dilutions were prepared with DMEM containing polybrene (EMD Millipore) to a final concentration of 8 µg/ml. Each well was infected with 75 µl of inoculum and incubated at 37 °C for 2 h rocking and rotating the plates every 10 min. After the 2 h adsorption, 250 µl of 5 % FBS/DMEM was added to each well. Twenty-four hours after infection, EGFP foci were visualized with a Nikon Eclipse TS2R microscope.

### Lentiviral infection and cell line production

Five hundred thousand HeLa cells were seeded into 100 mm dishes. About 18 h later, cells were infected at an estimated MOI of about 1 for each lentivirus in the presence of 8 µg/ml polybrene (EMD Millipore) in 1.5 ml inoculum and incubated at 37 °C for 2 h rocking and rotating every 10 min before adding 8.5 ml of 5 % FBS/DMEM. Medium was changed 24 h later to 10 ml of 5 % FBS/DMEM and changed again 48 h later to 10 ml of 5 % FBS/DMEM containing 1 µg/ml of puromycin (Mirus). Cells were incubated in selection medium for 72 h, when medium was replaced with 10 ml of 5 % FBS/DMEM containing 0.25 µg/ml of puromycin (maintenance medium).

### Transfections and fluorescence recovery after photobleaching (FRAP)

Transfections and FRAP were performed as described (31, 89–91). Nuclei of cells that were not attached to the coverslip, undergoing apoptosis or mitosis, that had blebbing or broken nuclear membranes, with too low fluorescence intensity (set as gain 600), or with punctate GFP-fluorescence (indicative of misfolded protein) were not included in the analyses. The dynamics of GFP-H2A were evaluated by FRAP approximately 48 hours after transfection, to ensure its assembly in chromatin during two S phases, whereas those of GFP-H2A.B, -H2A.X or -macroH2A were evaluated as early as 20 hours after transcription, as their assembly requires no S phase.

### Statistical analysis

Statistical significance was tested by one-tailed Student’s T test (for two treatments) or ANOVA with Tukey’s test post hoc (for more than two treatments).

### Immunofluorescence (IF)

IF was performed as described previously (90), with the following modifications. Two hundred thousand HeLa cells expressing flag tagged H2A, H2A.B, macroH2A, or H2B, were seeded onto sterile glass coverslips (18 mm diameter, 1.5 thickness). Cells were fixed in 4 % buffered formaldehyde (Sigma, F8775-25ml) freshly diluted in PBS, for 10 min at room temperature, permeabilized with 1 ml of 0.2 % Triton X-100 (Sigma) in PBS for 10 min at room temperature and blocked with 500 µl of 1 % bovine serum albumin ([BSA], Thermo Scientific), 5 % Goat Serum ([GS], Gibco), and 0.1 % Triton X-100 in PBS for 2 h at room temperature. Cells were incubated with 1:400 rabbit polyclonal anti-flag (ab205606, Abcam) in 0.1 % BSA, 1 % GS, 0.1 % Triton X-100 in PBS on a rocker at 4 °C overnight. Cells were incubated with 1:2000 Alexa Fluor 488 conjugated goat anti-rabbit (Invitrogen) in 0.1 % BSA, 1 % GS in PBS for 1 h at room temperature on a rocker, protected from light. Nuclei were counterstained with 1 µg/ml 4’,6-Diamidine-2’-phenylindole dihydrochloride (DAPI). Coverslips were mounted with 10 µl of ProLong^TM^ Gold antifade reagent (Invitrogen) after rinsing in deionized H_2_O. Image acquisition was performed with an Olympus FluoView FV3000 Confocal laser scanning microscope.

### Click chemistry labelling and detection of viral DNA

Wild-type or flag-tagged HeLa cell lines were infected with HSV-1 for 8 hours (MOI = 3). At 6hpi, 2.5μM 5-ethynyl-2’-deoxyuridine (Thermo Fisher A10044 or component of Click-iT® Alexa Fluor 488 reaction kit, C10337) and 2.5μM 5-ethynyl-2’ deoxycytidine (Sigma Aldrich T511307) nucleoside analogues were mixed in DMEM supplemented with 5% FBS media. One milliliter labelling solution was added to wells already containing one milliliter of media, for a final combined label concentration of 2.5μM (1.25μM each nucleoside). Click chemistry stock solutions and reaction buffer (Thermo Fisher Click-iT® Alexa Fluor 488 reaction kit, C10337) were prepared according to the manufacturer’s instructions right before using. After permeabilization, cells were washed twice with one milliliter of 3% BSA in PBS and incubated at room temperature for 30 minutes in 0.5 mL Click-iT® reaction cocktail with gentle rocking, protected from light. Cells were then incubated with blocking solution, primary antibodies, and secondary antibodies as described above using 1:400 rabbit polyclonal anti-flag primary antibody (ab205606, Abcam) or 1:400 mouse monoclonal ICP8 primary antibody (Abcam ab20194), and 1:2000 Alexa Fluor 546 conjugated goat anti-rabbit secondary antibody (Invitrogen). Remaining immunofluorescence steps were performed as previously described.

### Cell doubling

Puromycin was withdrawn from H2A-3XF, H2B-3XF, H2A.B-3XF, macro-H2A-3XF, and GFP HeLa cells. Forty-eight hours later, cells lines were seeded onto 96-well plates at 5.3 x 10^3^ cells per well. ATP levels were evaluated using the Promega CellTiter-Glo kit at 0, 24, and 48 hours post attachment (approximately 4 hours after seeding) following the manufacturer’s instructions.

### Viral growth curve

Puromycin was withdrawn from H2A-3XF, H2B-3XF, H2A.B-3XF, macro-H2A-3XF, and EGFP HeLa cells. Thirty-six hours later, cells were seeded onto 12-well plates at 1.5 x 10^5^ cells per well. Cells were infected with 100 µl HSV-1 KOS (MOI, 3.5 or 5.0). Inoculum was removed one hour later, unattached virions were washed away, and cells were overlaid with 1.0 ml 5 % FBS DMEM. One hundred microliters of supernatant were removed and replaced with 100 µl 5 % FBS DMEM at 3, 6, 9, 12, 15, 18, 30, and 36 hpi. Supernatants were immediately placed on ice and spun down at 500 g for 5 minutes at 4 °C. Eighty microliters of supernatant were transferred to a new tube, quickly frozen, and stored at -80 °C.

To evaluate viral titers, Vero-76 cells were seeded onto 12-well plates at a density of 1.4 x 10^5^ cells per well and infected with 10-fold serial supernatant dilutions. Inocula were removed one hour later, unattached virions were washed away, and infected cells were overlaid with 1 ml 5 % FBS 2 % methylcellulose DMEM. Cells were stained overnight with 1 ml crystal violet in 17 % methanol once plaques were visible and well-resolved. Viral titers were calculated from plaque counts.

### Standard chromatin immunoprecipitation (ChIP)

ChIP was performed as described previously (2), with the following modifications. Infected HeLa cells expressing H2A-3Xflag, H2B-3Xflag, H2A.B-3Xflag, or macroH2A-3Xflag (6 x 10^6^ on 100 mm dish) were cross-linked with 1 % of methanol-free formaldehyde (Thermo Scientific, 28906) in DMEM for 10 min at 37 °C. Chromatin was harvested and sheared as described (2). Sheared chromatin was thawed on ice and diluted with ChIP binding buffer (1 % Triton X-100, 10 mM Tris pH 8.0, 150 mM NaCl, 2 mM EDTA pH 8.0) to a final concentration of 0.58 ng/µl or 0.42 ng/µl for flag or H2B IP, respectively, to reach 350 or 250 ng of chromatin per IP reaction. One and a half microgram anti-flag antibody (Sigma-Aldrich, F1804), nonspecific mouse IgG (Sigma-Aldrich, M5284), anti-H2B antibody (Abcam, ab1790), or nonspecific rabbit IgG (Abcam, ab171870) per ChIP reaction, were conjugated to 0.225 mg of protein G Dynabeads (Invitrogen, 10003D), or protein A Dynabeads (Invitrogen, 10002D), for mouse or rabbit antibodies, respectively, for 2 h at RT with constant rotation. Three hundred and fifty or two hundred and fifty nanograms of chromatin for IP with mouse or rabbit antibodies, respectively, was incubated with bead-Ab complexes for about 20 h at 4 ° C with constant rotation. Immunopurified complexes were rinsed consecutively with 1.0 ml of each washing solution 1 (1 % Triton X-100, 2 mM EDTA, 20 mM Tris [pH 8.0], 150 mM NaCl), 2 (1 % Triton X-100, 2 mM EDTA, 20 mM Tris [pH 8.0], 500 mM NaCl) and 3 (1 % NP-40, 1 % NaDOC, 1 mM EDTA, 10 mM Tris [pH 8.0], 250 mM LiCl), were eluted with 600 µl of ChIP elution buffer (10 mM Tris-HCl [pH 8.0], 150 mM NaCl, 1 mM EDTA, 0.5 % SDS). Six microliters of proteinase K (20 mg/ml) (NEB, P8107S) were added, and samples were incubated at 65 ° C for approximately 19 h for de-crosslinking and deproteinization. Immunoprecipitated DNA was extracted with phenol-chloroform, isopropanol precipitated, ethanol washed, and subjected to qPCR (Fast SYBR Green master mix, Applied Biosystems, 4385612) with 300 nM each of GAPDH or UL46 primers using a QuantStudio3 from Applied Biosystems (Table 6). qPCR reactions were incubated 95 ° C for 20 sec denaturation, followed by 40 cycles of 3 sec denaturation at 95 ° C and 30 sec annealing/elongation at 65 ° C. Amplicon melting temperature was analyzed after the last cycle by 15 sec denaturation at 95 ° C and 60 sec annealing at 60 ° C, followed by a temperature ramp from 60 to 95 ° C in 0.1 ° C/sec increments and a final 15 sec incubation at 95 ° C.

Viral copy number was calculated from a standard curve using tenfold dilutions of 3,000,000 genome copy equivalents of cell free HSV-1 DNA spiked into 2.8 ng of HeLa cell DNA and sheared by sonication. Immunoprecipitation efficiencies were calculated by subtracting the genome copy equivalents co-immunoprecipitated with the non-specific IgG from those specifically precipitated with anti-flag or anti-H2B antibodies and are expressed as percentage of total genome copy equivalents in the harvested chromatin.

### Chromatin immunoprecipitation (ChIP) of serially MCN digested chromatin

ChIP was performed as described previously (2), with the following modifications. Infected HeLa cells expressing H2A-3Xflag, H2B-3Xflag, H2A.B-3Xflag, or macroH2A-3Xflag (6 x 10^6^ on 100 mm dish) were washed with 4 °C PBS, trypsinized, and resuspended in 10 ml of DMEM-5 % FBS. Cells were then pelleted at 500 x g for 5 min at 4°C and resuspended in 10 ml 4 °C PBS. Cells were pelleted at 500 x g for 5 min at 4°C and resuspended in 10 ml of 4 °C reticulocyte standard buffer (RSB,10 mM Tris [pH7.5], 10 mM NaCl, 5 mM MgCl2). Cells were pelleted at 500 x g for 5 min at 4°C and resuspended in 10 ml 4 °C RSB supplemented with protease inhibitor (cOmplete^tm^, EDTA-free protease inhibitor cocktail, Roche, Cat#: 11873580001) and incubated on ice for 15min. The cell membrane was lysed by adding 10 ml of RSB supplemented with 1 % (vol/vol) triton X-100 and protease inhibitor, to every 10 ml of cells in RSB and incubating on ice for 10 min, flipping the tubes upside down thrice. Nuclei were then pelleted by centrifugation at 3,250 x g for 25 min at 4 °C and transferred to 1.5 ml Eppendorf tubes using 500 μl of RSB supplemented with protease inhibitor. Nuclei were then pelleted by centrifugation at 10,000 x g for 1 min at 4 °C. Each 6 x 10^6^ nuclei were resuspended 1 ml chromatin extraction buffer (0.5 mM ethylene glycol-bis (beta-amino ethyl ether) -N, N, N’, N’, -tetra-acetic acid EGTA), 2 % triton X-100, 3 mM MgCl2, 20 mM Tris [pH8], 50 mM NaCl, 1 mM EDTA). Ten percent of the chromatin suspension was transferred into a new tube as non-digested and non-fractionated chromatin and kept on ice until the cross-linking step. Resuspended nuclei were then incubated for 1 h at 4 ° C with constant rotation. At the 30 min incubation time point, nuclei were mixed by pipetting up and down 10 times using a P1000 at a setting of 750 μl. Chromatin was spun down at 4,000 x g for 10 min at 4 °C and resuspended in 500 μl of ice cold MCN digestion buffer (10 mM Tris [pH8], 1 mM CaCl2). Chromatin was spun down again at 4,000 x g for 10 min at 4 °C and pelleted chromatin was disrupted by racking on a rack, resuspended in 110 μl per 6 x 10^6^ cells of MCN digestion buffer at room temperature. Ten microliter of MCN (0.005 U) were added to each sample with an offset of 20 seconds, each tube was flicked 10 times with the index finger to mix and spun down at 800 x g for 5 min. The supernatant (soluble chromatin) was removed with an offset of 20 seconds and transferred into previously prepared tube containing 90 μl of 0.05M EGTA on ice for immediate quenching of the MCN to prevent further digestion of the DNA released in the soluble chromatin. The pellet (insoluble chromatin) was disrupted as before and resuspended with fresh 110 μl of MCN digestion buffer and placed on ice until the last sample is collected. The entire procedure was repeated six times. After the sixth round of digestion, 810 µl of MCN buffer with 0.005M EGTA were immediately added to quench MCN to the pellet and to wash any soluble fragments left behind. The pellet was then resuspended with 810 µl of MCN buffer with 0.005M EGTA and kept on ice until the cross-linking stage. The supernatants from the six rounds of digestion were pooled and centrifuged at 15,000 x g at 4 °C for 20 min. The supernatant was then transferred to a new tube (Short-soluble fraction). The pelleted fraction from the supernatant was resuspended with 810 µl of MCN buffer with 0.005M EGTA (Long-soluble fraction) and placed on ice until the cross-linking stage.

The samples containing short-soluble, long-soluble, insoluble, and undigested chromatin were adjusted to 810μl with MCN-digestion buffer with 0.005M EGTA and cross-linked by adding 135 μl of buffered 7 % formaldehyde solution (7 % CH2O, 140 mM Na2HPO4 pH 8.67) using 16 % CH_2_O methanol-free ampules (Thermo Scientific Cat #28906), to a final concentration of 1 %, for 10 min at 4 °C with constant mixing. Cross-linking was quenched with 63 μl of 2 M glycine, to a final concentration of 125 mM, for 5 min at room temperature. Ten percent of each sample was transferred into new tubes for DNA purification and digestion check.

The samples containing long-soluble, insoluble, and undigested chromatin were sonicated with a Diagenode Bioruptor Plus in a 4 °C recirculating water bath. The undigested and insoluble chromatin were sonicated for 120 min with the power setting on high and with 30 seconds of pulse on followed by 30 seconds of pulse of for total of 60 min sonication time. Ten percent of each sample was transferred into new tubes for DNA purification and quantitation. The remaining samples were flash frozen with liquid nitrogen and stored at – 80 °C until the chromatin immunoprecipitation step.

Sheared and digested chromatin was thawed on ice and diluted with ChIP binding buffer (1 % Triton X-100, 10 mM Tris pH 8.0, 150 mM NaCl, 2 mM EDTA pH 8.0) to a final concentration of 0.17 ng/µL for flag or H2B IP, to reach 100 ng of chromatin per IP reaction. One and a half microgram anti-flag antibody (Sigma-Aldrich, F1804), nonspecific mouse IgG (Sigma-Aldrich, M5284), anti-H2B antibody (Abcam, ab1790), or nonspecific rabbit IgG (Abcam, ab171870) per ChIP reaction, were conjugated to 0.225 mg of protein G Dynabeads (Invitrogen, 10003D), or protein A Dynabeads (Invitrogen, 10002D), for mouse or rabbit antibodies, respectively, for 2 h at RT with constant rotation. One hundred nanograms of chromatin for IP with mouse or rabbit antibodies was incubated with bead-Ab complexes for about 20 h at 4 ° C with constant rotation. Immunopurified complexes were rinsed consecutively with 1.0 ml of each washing solution 1 (1 % Triton X-100, 2 mM EDTA, 20 mM Tris [pH 8.0], 150 mM NaCl), 2 (1 % Triton X-100, 2 mM EDTA, 20 mM Tris [pH 8.0], 500 mM NaCl) and 3 (1 % NP-40, 1 % NaDOC, 1 mM EDTA, 10 mM Tris [pH 8.0], 250 mM LiCl), were eluted with 600 µl of ChIP elution buffer (10 mM Tris-HCl [pH 8.0], 150 mM NaCl, 1 mM EDTA, 0.5 % SDS). Six microliters of proteinase K (20 mg/ml) (NEB, P8107S) were added, and samples were incubated at 65 ° C for approximately 19 h for de-crosslinking and deproteinization. Immunoprecipitated DNA was extracted with phenol-chloroform, isopropanol precipitated, ethanol washed, and subjected to DNA sequencing library preparation or analyzed through qPCR (Fast SYBR Green master mix, Applied Biosystems, 4385612) with 300 nM each of GAPDH or UL46 primers using a QuantStudio3 from Applied Biosystems (Table 6). qPCR reactions were incubated 95 ° C for 20 sec denaturation, followed by 40 cycles of 3 sec denaturation at 95 ° C and 30 sec annealing/elongation at 65 ° C. Amplicon melting temperature was analyzed after the last cycle by 15 sec denaturation at 95 ° C and 60 sec annealing at 60 ° C, followed by a temperature ramp from 60 to 95 ° C in 0.1 ° C/sec increments and a final 15 sec incubation at 95 ° C. Immunoprecipitation efficiencies were calculated as before.

### DNA sequencing library preparation

Total DNA was quantitated using a Qubit 4 Fluorometer (Thermo Fisher) and following the manufacturers protocol. Sequencing libraries were constructed using 1.34 ng of input DNA and following the manufacturers protocol of NEBNext Ultra^tm^ II DNA library prep kit for Illumina. Final libraries were analyzed using a Qubit Fluorimeter and submitted to Cornell Institute of Biotechnology for sizing and quality assessment with a fragment analyzer (AATI, Agilent) prior to pooling for sequencing. The area under the curve of the DNA library fragment analyzer plots was used to calculate the average molarity of the DNA fragments from the range of 200 to 700 bp. Individual libraries were pooled at equimolar concentrations and resolved in a 1.5 % agarose gel at 70 V for 1.5 h. DNA fragments from 200 to 700 bp were gel purified using a QIAquick gel extraction kit following the manufactures protocol. Library pools where libraries were analyzed using a Qubit Fluorimeter and submitted to Cornell Institute of Biotechnology for sizing and quality assessment with a fragment analyzer (AATI, Agilent) prior to sequencing. DNA sequencing libraries from untreated infections, traditional ChIP and ChIP from serially MCN digested and fractionated chromatin, were sequenced across 4 HiSeq PE150 lanes. DNA sequencing libraries from CHX treated infections were sequenced across 1 NovaSeq PE150 lane. Sequencing was performed by Novogene.

### Sequencing analysis

Demultiplexed datasets provided by Novogene were trimmed to remove adaptor sequences and low quality 3’ nucleotides with TrimGalore [ --paired --length 30 –q 30 ] (https://www.bioinformatics.babraham.ac.uk/projects/trim_galore/). Resulting reads were aligned against a combined human (hg38) and HSV-1 (strain KOS, KT899744) genome reference using bowtie2 (116) and subsequently processed with SAMtools (117) and BEDtools (118) to separate the alignments. Sambamba (119) was to filter out poor quality alignments [ -F not (unmapped or mate_is_unmapped or supplementary or secondary_alignment) and proper_pair ], while reformat.sh from the BBmap (https://sourceforge.net/projects/bbmap/) package was used to randomly subsample 50,000 paired reads from each HSV-1 alignment. For some samples, this subsampling was reduced to 12,000 paired reads due to limited numbers of HSV-1 aligning reads. LSBED4 coverage files were generated from the randomly subsampled alignments with genomeCoverageBed (BEDTools). The detailed account of the sequencing reads is provided in Supplementary Table 1.

### Plotting

All plotting was performed using R studio (120) with R v4.1.1 and the following packages: data.table, Gviz (121), Genomic Features (122). Briefly, paired BED4 files (input & IP for a given condition/antibody) were imported and the depth of coverage at each position expressed as log2(IP_depth/input_depth). Genome wide plots were generated using Gviz and annotated with the strain KOS v1.2 annotation (Depledge, pers comm).

### Data availability

All Illumina sequencing datasets associated with this study are available as raw paired fastq files via the European Nucleotide Archive under the accession PRJEB65576. Full details regarding each sequencing dataset are in Supplementary Table 1 All R scripts and underlying datasets used to generate analyses and associated visualizations of the sequencing data are available at (https://github.com/DepledgeLab/HSV1-chromatin-dynamics).

## Acknowledgments

These studies were funded by the National Institutes of Health (NIH) IAID 1R01AI153396, and previously by the Canadian Institutes of Health Research. We acknowledge the support from Cornell College of Veterinary Medicine and the Baker Institute for Animal Health. LMS is a BWF investigator in the Pathogenesis of Infectious Disease. EF acknowledges the support from the Consejo Nacional de Ciencia y Tecnologia (CONACyT) and the Biomedical and Biological Science Field (Cornell University).

## Supplementary Material

**Figures S1, S2**

**Table S1.** Summary information of the sequencing data

**Figure S1. Endogenous histone H2B and flag tagged histone H2A, H2A.B, macroH2A, or H2B are homogeneously distributed throughout the HSV-1 genome regardless of the dynamic state of the HSV-1 chromatin.** Line graphs presenting the log2 HSV-1 coverage plots from 50,000 HSV-1-aligned paired-end reads normalized IP coverage (Flag IP/No IP) or (H2B IP/ No IP); Sh, short accessible chromatin; Lo, long accessible chromatin; Pe, insoluble chromatin; ND NF, non-digested non-fractionated but sonicated chromatin; Gray rectangles, cartoon representation of the HSV-1 genome.

**Figure S2. Flag tagged histones are equally incorporated into chromatin along the entire HSV-1 genome regardless of whether the entire genome is transcribed or transcription is restricted to the IE loci.** Genome coverage of the normalized signal from the co-immunoprecipitation of each flag tagged histone divided by the normalized H2B signal in infections treated or not with CHX.

